# Systematic Establishment of Robustness and Standards in Patient-Derived Xenograft Experiments and Analysis

**DOI:** 10.1101/790246

**Authors:** Yvonne A. Evrard, Anuj Srivastava, Jelena Randjelovic, Sasi Arunachalam, Carol J. Bult, Huiqin Chen, Lily Chen, Michael Davies, Sherri Davies, Brandi Davis-Dusenbery, Jack DiGiovanna, Li Ding, James H. Doroshow, Bingliang Fang, Christian Frech, Ramaswamy Govindan, Min Jin Ha, Meenhard Herlyn, Ryan Jeon, Andrew Kossenkov, Michael T. Lewis, Shunqiang Li, Michael Lloyd, Funda Meric-Bernstam, Nevena Miletic, Jeffrey A. Moscow, Steven Neuhauser, David Nix, Rajesh Patidar, Vito Rebecca, Peter N. Robinson, Jacqueline Rosains, Jack Roth, Isheeta Seth, Tamara Stankovic, Adam Stanojevic, Brian A. Van Tine, Alana L. Welm, Bryan E. Welm, Jayamanna Wickramasinghe, XingYi Woo, Min Xiao, Zi-ming Zhao, Dennis A. Dean, Jeffrey S. Morris, Jeffrey H. Chuang

## Abstract

Patient-Derived Xenografts (PDXs) are tumor-in-mouse models for cancer. PDX collections, such as those supported by the NCI PDXNet program, are powerful resources for preclinical therapeutic testing. However, variations in experimental design and analysis procedures have limited interpretability. To determine the robustness of PDX studies, the PDXNet tested temozolomide drug response for three pre-validated PDX models (sensitive, resistant, and intermediate) across four blinded PDX Development and Trial Centers (PDTCs) using independently selected SOPs. Each PDTC was able to correctly identify the sensitive, resistant, and intermediate models, and statistical evaluations were concordant across all groups. We also developed and benchmarked optimized PDX informatics pipelines, and these yielded robust assessments across xenograft biological replicates. These studies show that PDX drug responses and sequence results are reproducible across diverse experimental protocols. Here we share the range of experimental procedures that maintained robustness, as well as standardized cloud-based workflows for PDX exome-seq and RNA-Seq analysis and for evaluating growth.

## Introduction

Patient-Derived Xenografts (PDX) are in vivo preclinical models in which human cancers are engrafted into a mouse for translational cancer research and personalized therapeutic selection ^1–8^. Prior studies have shown that treatment responses of tumor-bearing mice usually reflect the responses in patients ^5,9,10^. PDXs have been used successfully for preclinical drug screens ^5,8,9^, to facilitate the identification of potential biomarkers of drug response and resistance ^5,8,11,12^, to select appropriate therapeutic regimens for individual patients ^13^, and to measure evolutionary processes in cancer in response to treatment ^14^. At the genomic level, engrafted human tumors have been shown to retain most genomic aberrations from the original patient tumor ^13,15,16^. These successes have led to the development of a number of PDX collections in both academia and industry ^9,17–19^ for use in preclinical testing.

Despite these successes, important questions remain for the use of PDXs as a model system for treatment response. The reproducibility of treatment response has not been well-evaluated because research teams frequently do not repeat large in vivo experiments, and often perform experiments in models that are not used by other groups. Variations in engraftment rates for different cancer subtypes, study duration, dosing routes and schedules, and response assessment protocols also frustrate comparisons of results. Moreover, intratumoral heterogeneity within primary tumors, as well as potential genetic drift or selection (“bottlenecking”) during tumor engraftment and xenograft passaging, can result in genomic variation among primary tumor samples and their subsequently-derived xenografts ^5,16^. Whether such variation impacts the accuracy of PDXs as a preclinical model has been unclear. Resolution of this issue requires comparison of primary tumors and their derivative PDX to ensure biological consistency whenever possible, as well as controlled treatment regimens with human-equivalent dosing.

Resolution of this issue at the molecular level will also require standardized PDX-specific sequence analysis pipelines to robustly identify genomic aberrations either selected for, or arising de novo, in the PDX model relative to the tumor of origin. Progress on these topics is important to the overall field of cancer patient-derived models, as analogous concerns pertain to organoids and other 3D culture systems.

To resolve such questions related to the use of PDXs in precision medicine, the US National Cancer Institute has supported a consortium of PDX-focused research enters, the NCI PDXNet. Here, we in the PDXNet consortium report the results of experiments to test the robustness of PDX treatment responses across different research centers, using temozolomide treatment on three blinded models chosen by the NCI Patient Derived Models Repository based on existence of prior data on their temozolomide responses. We report on replicate evaluations across four Patient-derive Xenograft Development and Trials Centers (PDTC) of the PDXNet using blinded treatment and response evaluation protocols uniquely chosen by each PDTC.

Simultaneously, we performed exome and RNA sequencing at each center to determine biological and technical stability of genomic characterizations of samples from each center. These sequence analyses were performed with optimized analysis pipelines chosen based on an extensive new benchmarking of pipelines from each center on synthetic sequence sets.

Finally, we have statistically analyzed the cohort growth curves for each model in each research center using five separate metrics. These studies allowed us to assess whether PDXs have sufficiently robust behaviors to withstand variations in experimental procedures, response measurement algorithms, genomic variation among replicates, and alternative sequence analysis protocols. We also report effective SOPs for experimental procedures, pipelines for statistical assessment of response, as well as both DNA and RNA sequence analysis workflows. We expect these standards to advance the use of PDXs and other in vivo models in cancer precision medicine, a critical need for the evaluation of PDX results in the context of moving novel therapeutics or therapeutic combinations to the clinic.

## Results

### Study design and treatment results

A critical, yet unresolved, question that motivated the inception of the PDXNet was what the inter-laboratory reproducibility of PDX drug studies would be across centers with independently established practices for preclinical testing, i.e. how much standardization would be needed to run large-scale, multicenter preclinical studies. To address this question, the NCI Patient Derived Models Repository (PDMR) reviewed preclinical studies performed by the Biological Testing Branch (BTB/DCTD/NCI), which has performed numerous in vivo studies with PDX models. The PDMR selected three PDX models with non-published known responses to temozolomide for an inter-laboratory reproducibility pilot. The three PDX models selected were 625472-104-R (colon adenocarcinoma, non-responsive model), 172845-121-T (colon adenocarcinoma, intermediate response), and BL0293-F563 (urothelial/bladder cancer, complete response). Patient data, clinical history, and representative histology and sequence data can be found at https://pdmr.cancer.gov; model BL0293-F563 was originally developed by The Jackson Laboratory (tumor model TM00016, http://tumor.informatics.jax.org/mtbwi/pdxSearch.do).

For the study set-up (Figure 1), the four PDTCs--Huntsman Cancer Institute/Baylor College of Medicine (HCI-BCM), MD Anderson Cancer Center (MDACC), Washington University-St. Louis (WUSTL), and The Wistar Institute/University of Pennsylvania/MDACC (WIST) were directed to use their standard preclinical study set-up and monitoring SOPs and to use literature searches to determine temozolomide dosing and schedule. All PDTCs were kept blinded to which models were temozolomide sensitive or resistant and to all other groups’ preclinical study set-ups. In addition, none of the PDTCs had previous experience with temozolomide; so the reference doses/schedules would need to be determined independently at each center. The exceptions to blinding were that all PDTCs were required to use NSG host mice and implant PDX material subcutaneously. In addition, the PDTCs used drug prepared by the Developmental Therapeutics Clinic (DTP/NCI) to ensure that there were no variations in manufacture.

**Figure 1.**
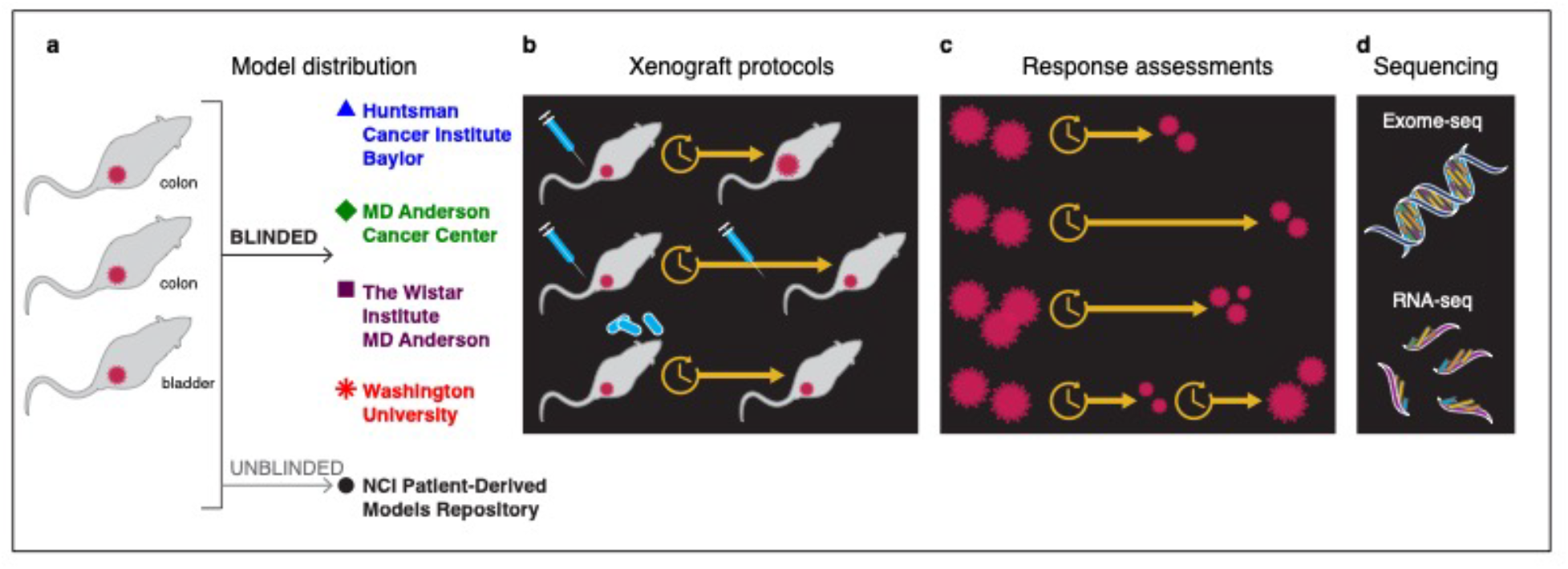
**a)** Three models were distributed for experimentation to 4 centers: MD Anderson, WUSTL, Wistar, and HCI/BCM. These three centers were chosen based on prior results on temozolomide treatment response obtained by the PDMR. **b)** Each of the three models were treated with temozolomide by the 4 centers under blinded protocols. **c)** Treatment responses were comparatively assessed under several biostatistical protocols. **d)** Sequence data were collected by each center and assessed.

The laboratory SOPs for the preclinical study set-ups were collated by the PDMR (Table 1). While all centers staged tumors to between 100-200 mm^3^, implantation methodologies varied. Three groups directly implanted ~1 mm^3^ PDX fragments into each host mouse, one group minced a ~1 mm^3^ PDX fragment into a slurry for implantation, and one dissociated PDX material and implanted 3-5 × 10^6^ cells per host. Comparison of vehicle control growth curves for all groups demonstrated overall similar growth kinetics of the models at each site irrespective of the implantation methodology used (**Supplementary Figure 1**).

**Table 1.**
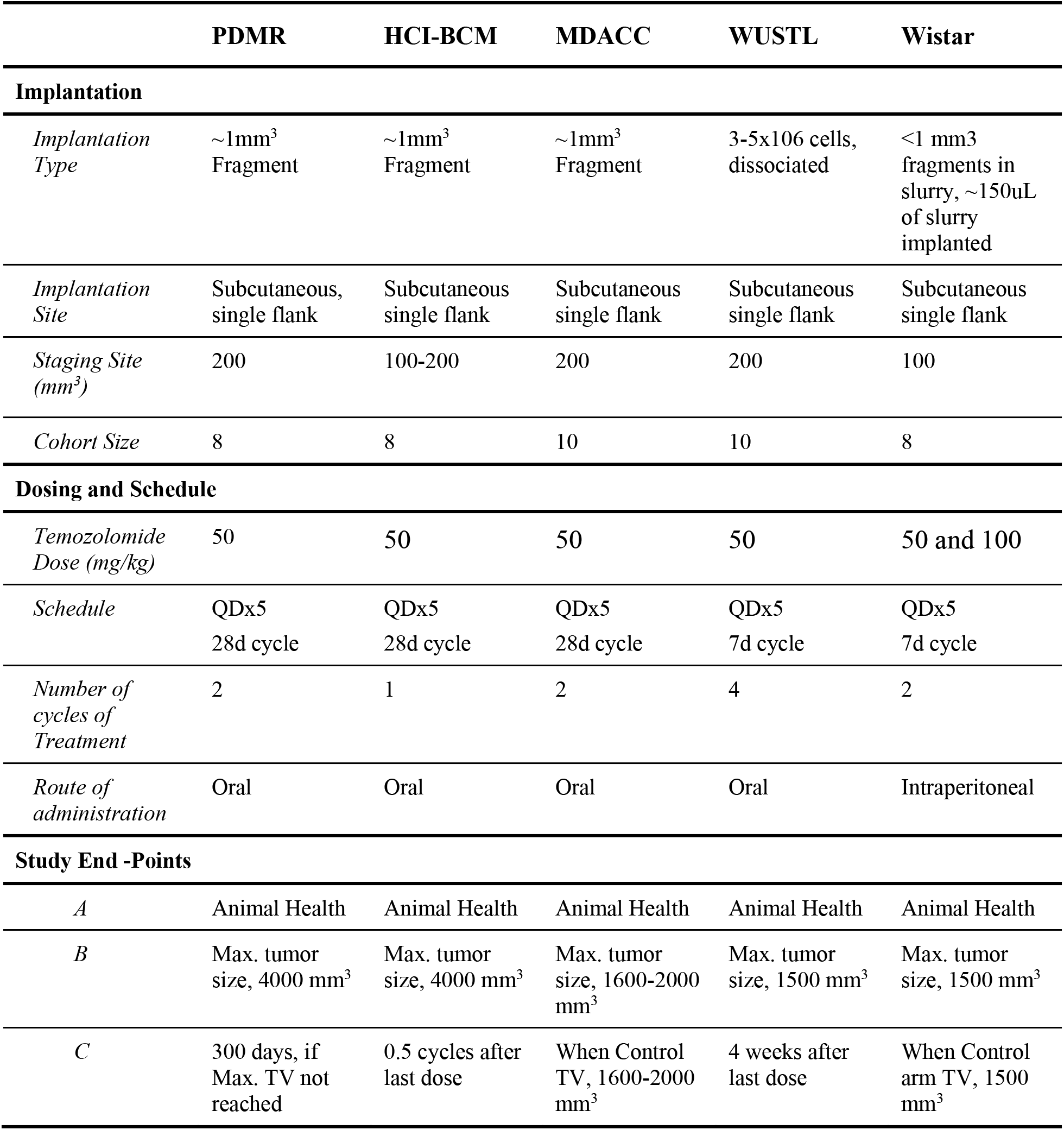
Comparison of preclinical study set-ups and end-points at the PDMR and individual PDTCs for the temozolomide Reproducibility Pilot. Abbreviations: QDx5 (Once daily for 5 days), TV (tumor volume).

Each PDTC independently researched published literature to select a temozolomide dosing and schedule for its site, with key references noted: HCI-BCM ^20–22^, MDACC ^23^, WUSTL ^22,24–27^, and WISTAR ^28,29^. While diverse literature was considered, all sites selected a 50 mg/kg dose and one of two different dosing schedules. These schedules were either daily temozolomide treatment for 5 days followed by 23 days of rest (28 day cycle) or 5 days of treatment followed by 2 days of rest (7 day cycle); 1-4 cycles were used (Table 1).

Overall, all sites reported similar responses irrespective of the methodology, dosing, or schedule used (Figure 2), with especially strong concordance in the non-responsive and completely responsive model results. If the drug x model combination had been performed as part of an exploratory study, these independent experiments would likely yield similar decisions about treatment efficacy. The intermediate response models showed more variation in growth across centers. The intermediate cases were also more clearly affected by the variability in SOP end-point times, one of the biggest variations among methodologies (Table 1). For example, some groups sacrificed all mice once the vehicle control group reached a threshold volume, while other groups ended after a defined length of time after the last dosing. This resulted in some studies observing strong tumor inhibition through the end of study, while others observed regrowth after initial inhibition **(Supplementary Figure 2)**. Nevertheless, the similarities in response indicated that the existing range of methodologies is sufficient and robust enough to capture the critical cases of strong response and non-response. After discussion of these results, the PDXNet Consortium has agreed on a standard of continued monitoring of all cohorts where response is observed for at least 1.5-2 cycle lengths beyond the last dosing cycle, provided animal health end-points are not reached. Detailed quantitative comparisons are addressed in the next section.

**Figure 2.**
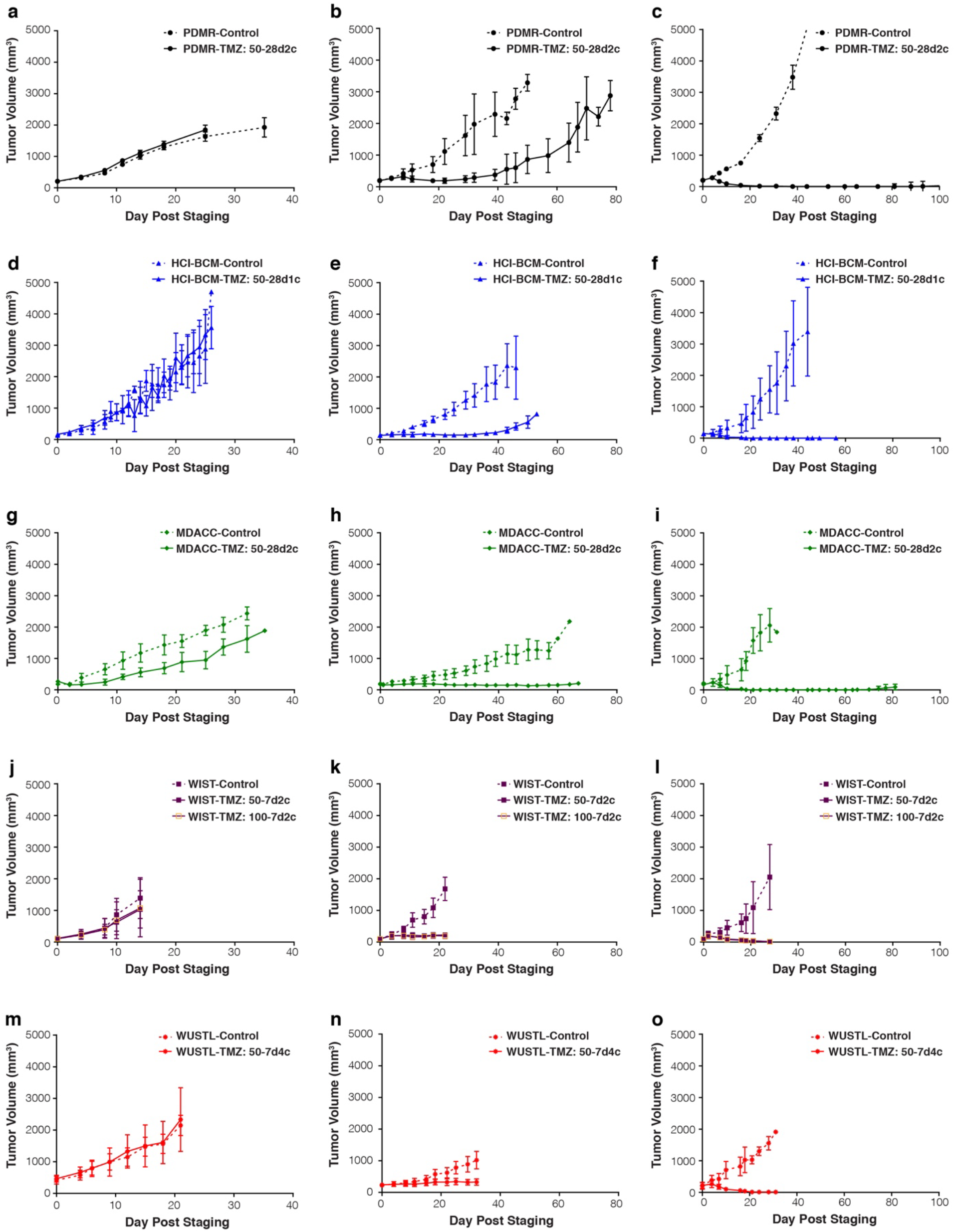
Comparison of PDX tumor volume control and temozolomide treatment arms at the PDMR **(a-c)**, HCI-BCM (d-f), MDACC **(g-i)**, WIST **(j-l)**, and WUSTL **(m-o)**. Model 625472-104-R **(a, d, g, j, m)**, 172845-121-T **(b, e, h, k, n)**, and BL0293-F563 **(c, f, i, l, o)**. Axes are held constant for comparison between studies. Dashed lines, vehicle control groups, Solid lines, temozolomide treatment groups. Median ± SD.

## Statistical Robustness of PDX Treatment Response

### Statistical approaches for evaluating cohort drug response

A challenge of evaluation of PDX response is that there is still no standard statistical approach for analysis of tumor response for PDX growth data. Common measures of tumor size include percent- or fold-change in volume from baseline to a fixed time end-point; area under the tumor growth curve; tumor growth inhibition, defined as the ratio of the average tumor size at a given time point relative to control; and time to event analysis, a potentially censored end-point measuring time from baseline until growth to a certain multiple of baseline. Classification of growing PDX tumors into RECIST-like categories ^30^ (Complete Response-CR, Partial Response-PR, Stable Disease-SD, and Progressive Disease-PD) is another assessment that has the advantage of congruence with clinical trials, but it can be strongly dependent on category thresholds that do not analogize straightforwardly with patient primary tumors. Each of these measures has their own strengths and limitations. For example, the percent change from baseline is intuitive, interpretable, and unlike RECIST-like criteria, does not require specification of a cut point. In contrast to the area under the curve (AUC) approaches percent- or fold change does not use all of the tumor time course information but only the first and last points. Here we consider all of these measures and assess concordance of results across analytical strategies as well as across growth data from each center.

### PDX tumor volume analysis software

We have devised an automated analysis script in R that, given data in a prespecified format and a time point of interest, will automatically plot the tumor growth curves and group mean curves, compute all of these statistical measures and their associated plots, and produce an annotated. html report in R markdown that serves as a complete summary of the results (see Methods). In the supplementary materials **(Supplementary Table 1)**, we describe a standard format for the recorded data that is compatible with our analysis scripts, and we also provide instructions for researchers to use this script to analyze their own data. We believe that this automated script can enhance reproducibility and transparency of analyses, and can be revised and adapted as a standard analysis script for general use.

### Comparisons across statistical methods

We statistically assessed drug response for the measures mentioned above across all research groups. Table 2 contains the p-values for assessing treatment vs. control differences for each of the statistical tests (see Methods), and Figure 3 shows associated plots for the three models from the HCI-BCM studies. Associated plots for drug response at other sites i.e. MD Anderson, Washington University, PDMR and Wistar are shown in **Supplementary Figures 3, 4, 5, and 6**, respectively. Overall, we found that assessments of drug response were robust across research groups, with particularly decisive evaluations for the non-responsive and responsive models. The various analytical methods **(Supplementary Figures 7, 8, 9 and 10)** also gave results consistent with one another, with a few exceptions noted below. However, the intermediate group was difficult to classify. For the intermediate group most of the statistical measures showed clear difference from control, but the results were inconsistent for RECIST-like criteria.

**Table 2.** Statistical tests of treatment vs. control difference Statistical tests of treatment vs. control difference. This table presents the p-values reported for various analytical measures, including change from baseline to 21 days (***ΔV*_21_**), adjusted area under the curve for 21 days (***αAUC*_21_**), adjusted area under the curve until last measurement (***αAUC*_*max*_**), tumor growth inhibition at day 21 (***TGI*_21_**), and RECIST-like criteria for various choices of boundaries between CR/PR, PR/SD, and SD/PD given by (*****c***_1_, ***c***_2_, ***c***_3_**). For RECIST-like, p-values are testing PD vs. not PD. * For the Wistar site, the first row is for TMZ 50mg/kg and the second row is TMZ 100 mg/kg.

**PD (Model 625472-104-R)**
| Site | $\Delta_{V_{21}}$ | $aAUC_{21}$ | $aAUC_{max}$ | $TGI_{21}$ | $RECIST_{21}$ | | | | | |
| --- | --- | --- | --- | --- | --- | --- | --- | --- | --- | --- |
|  |  |  |  |  | -95,-50,10 | -95,-30,20 | -95,-30,50 | -95,-30,100 | -95,-50,50 | -95,-50,100 |
| MDA | 0.163 | 0.236 | 0.448 | 0.067 | 1.000 | 1.000 | 1.000 | 1.000 | 1.000 | 1.000 |
| WU | 0.918 | 0.376 | 0.470 | 0.538 | 1.000 | 1.000 | 1.000 | 1.000 | 1.000 | 1.000 |
| HCI-BCM | 0.143 | 0.072 | 0.177 | 0.814 | 1.000 | 1.000 | 1.000 | 1.000 | 1.000 | 1.000 |
| PDMR | 0.404 | 0.756 | 0.501 | 0.751 | 1.000 | 1.000 | 1.000 | 1.000 | 1.000 | 1.000 |

**SD (Model 172845-121-T)**
| Site | $\Delta_{V_{21}}$ | $aAUC_{21}$ | $aAUC_{max}$ | $TGI_{21}$ | $RECIST_{21}$ | | | | | |
| --- | --- | --- | --- | --- | --- | --- | --- | --- | --- | --- |
|  |  |  |  |  | -95,-50,10 | -95,-30,20 | -95,-30,50 | -95,-30,100 | -95,-50,50 | -95,-50,100 |
| MDA | <.001 | 0.003 | <.001 | <.001 | 0.048 | 0.048 | 0.008 | 0.048 | 0.008 | 0.048 |
| WU | <.001 | <.001 | <.001 | <.001 | 1.000 | 0.474 | 0.211 | <.001 | 0.211 | <.001 |
| HCI-BCM | <.001 | <.001 | <.001 | <.001 | 0.200 | 0.026 | <.001 | <.001 | <.001 | <.001 |
| PDMR | <.001 | <.001 | <.001 | <.001 | 0.003 | <.001 | <.001 | <.001 | <.001 | <.001 |
| Wistar* | <.001 | <.001 | <.001 | <.001 | 1.000 | 1.000 | 1.000 | 0.200 | 1.000 | 0.200 |
|  | <.001 | <.001 | <.001 | <.001 | 1.000 | 1.000 | 0.200 | 0.026 | 0.200 | 0.026 |

**CR (Model BL0293-F563)**
| Site | $\Delta_{V_{21}}$ | $aAUC_{21}$ | $aAUC_{max}$ | $TGI_{21}$ | $RECIST_{21}$ | | | | | |
| --- | --- | --- | --- | --- | --- | --- | --- | --- | --- | --- |
|  |  |  |  |  | -95,-50,10 | -95,-30,20 | -95,-30,50 | -95,-30,100 | -95,-50,50 | -95,-50,100 |
| MDA | <.001 | <.001 | <.001 | <.001 | <.001 | 0.008 | 0.008 | 0.008 | 0.008 | 0.008 |
| WU | <.001 | <.001 | <.001 | <.001 | <.001 | <.001 | <.001 | <.001 | <.001 | <.001 |
| HCI-BCM | <.001 | <.001 | <.001 | <.001 | <.001 | <.001 | <.001 | <.001 | <.001 | <.001 |
| PDMR | <.001 | <.001 | <.001 | <.001 | <.001 | <.001 | <.001 | <.001 | <.001 | <.001 |
| Wistar* | <.001 | <.001 | <.001 | <.001 | <.001 | <.001 | <.001 | <.001 | <.001 | <.001 |
|  | <.001 | <.001 | <.001 | <.001 | <.001 | <.001 | <.001 | <.001 | <.001 | <.001 |

**Figure 3.**
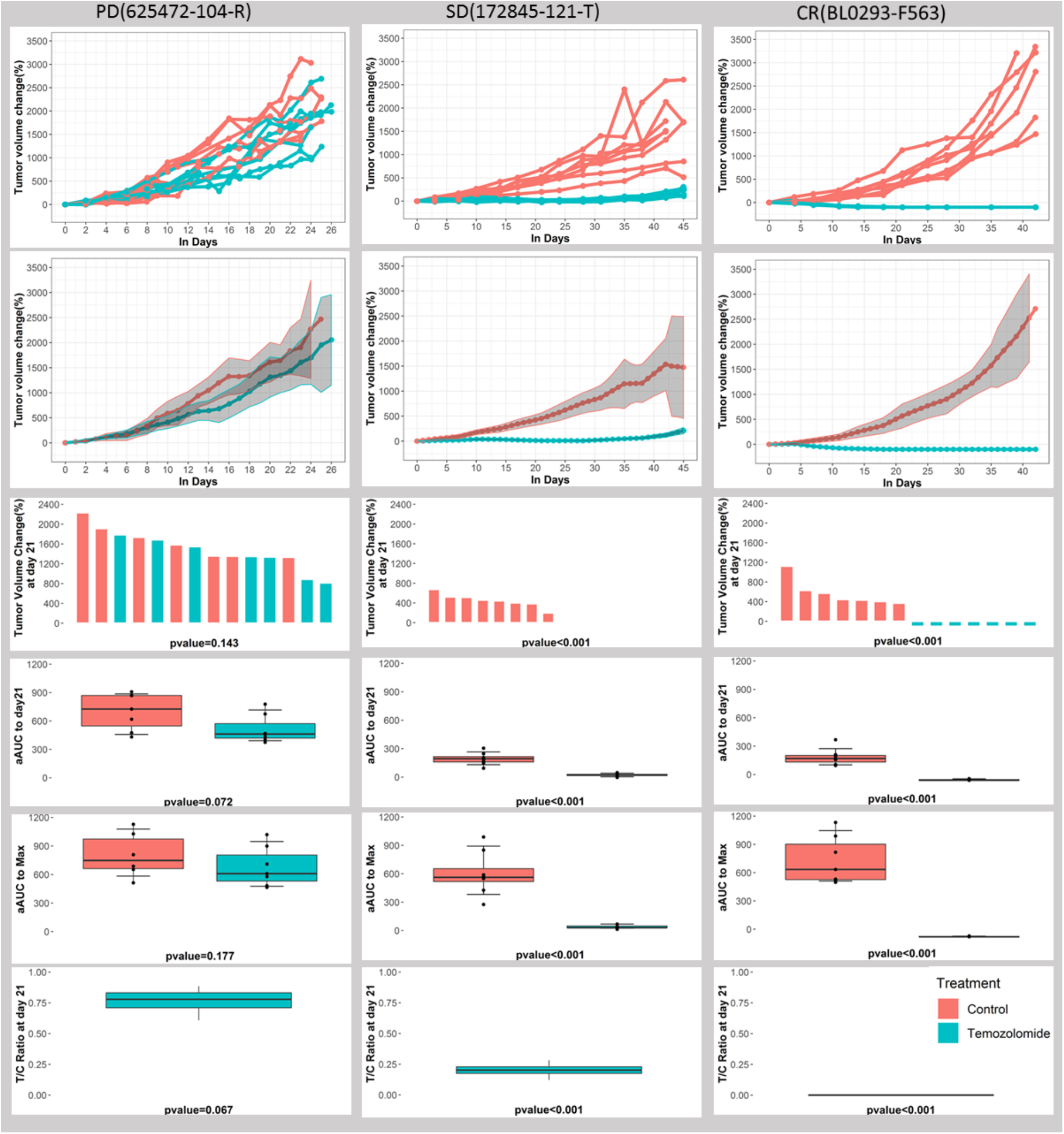
Analytical Summaries, HCI-BCM Study. Analytical results from the HCI-BCM study for progressive model (625472-104-R), stable disease model (172845-121-T) and complete response model (BL0293-F563) (columns 1-3, respectively), with interpolated individual curves (row 1), mean curves for treatment and control with 95% confidence bands (row 2), waterfall plots demonstrating ***ΔV*_21_**(row 3), boxplots of ***αAUC*_21_**(row 4) and ***αAUC*_*max*_**(row 5) for treatment and control, and a boxplot of ***TGI*_21_** (row 6), along with p-values comparing treatment to control for each measure.

RECIST yielded qualitatively similar ordering of the models as the other methods, but it had the lowest power and showed considerable variability across cut points, complicating its use. The percent change in tumor size and area under the curve measures largely agreed, and showed good statistical power. The tumor growth inhibition measure also yielded consistent results. The natural statistical test is whether this ratio is less than 1, but this should be accompanied by an assessment of the clinical significance of the effect size, since it is possible to have a small p-value with minimal inhibition, e.g. 10% or 20%, that might not be clinically meaningful. We recommend statistical testing vs. control while accompanied by an assessment of clinical significance that may depend on the context.

### Cloud Workflows for PDX Sequence Analysis

Robust sequence analysis pipelines are essential for understanding cancer genetics from PDX models. While prior PDX pipelines have been published, e.g. ^5,31^, it can be time-consuming for researchers to implement and evaluate other groups’ methods. To address this problem, five PDXNet teams provided sequence analysis workflows for PDX exome-seq mutation calling, and the PDXNet Data Commons and Coordinating Center (PDCCC) dockerized these for co-localized application and sharing with the research community via the National Cancer Institute Cancer Genomics Cloud (CGC). The Seven Bridges Genomics team in the PDCCC also independently evaluated each of these pipelines. Each submitting group also specified parameters as part of the workflow submission. Evaluations were performed on simulated benchmark mixtures of human and mouse reads with various mouse/human read ratios and variant allele frequencies (see Methods).

### Benchmarking of human-mouse read disambiguation

We first compared the efficacy of the five pipelines **(Supplementary Table 2)** for human-mouse read disambiguation using a series of simulated benchmark WES and RNA-Seq datasets. The WES benchmark dataset consisted of a paired end exome-seq data series with human-mouse ratios of 90:10, 80:20, 60:40, 50:50, 40:60, and 20:80 created by computationally mixing two 100% pure WES human and mouse samples (sample ids 14311X2 and 14311X8, respectively). Similarly, the RNA-Seq benchmark dataset consisted of a simulated human-mouse mixture series (90:10, 80:20, 60:40, 50:50, 40:60, 20:80) created from human and mouse lung RNA-seq samples (ENCSR129KCJ and ENCSR870AQU, respectively). The above-mentioned simulated WES and RNA-Seq datasets were used to test the five commonly used human-mouse read deconvolution tools: BBSplit, Xenome, Disambiguate, Xenofilter, and ICRG. All tools achieved >99 % precision for both WES and RNA-Seq benchmarks (Figure 4). Xenofilter showed the lowest recall (96.60 % and 89.63 % recall in WES and RNA-seq benchmarks, respectively), whereas BBSplit showed the best overall performance i.e. highest precision without any loss in recall (99.87 % and 99.64 % precision in WES and RNA-seq benchmarks, respectively), followed closely by Disambiguate on WES data (99.77 % precision) and Xenome on RNA-seq data (99.77 % precision).

**Figure 4.**
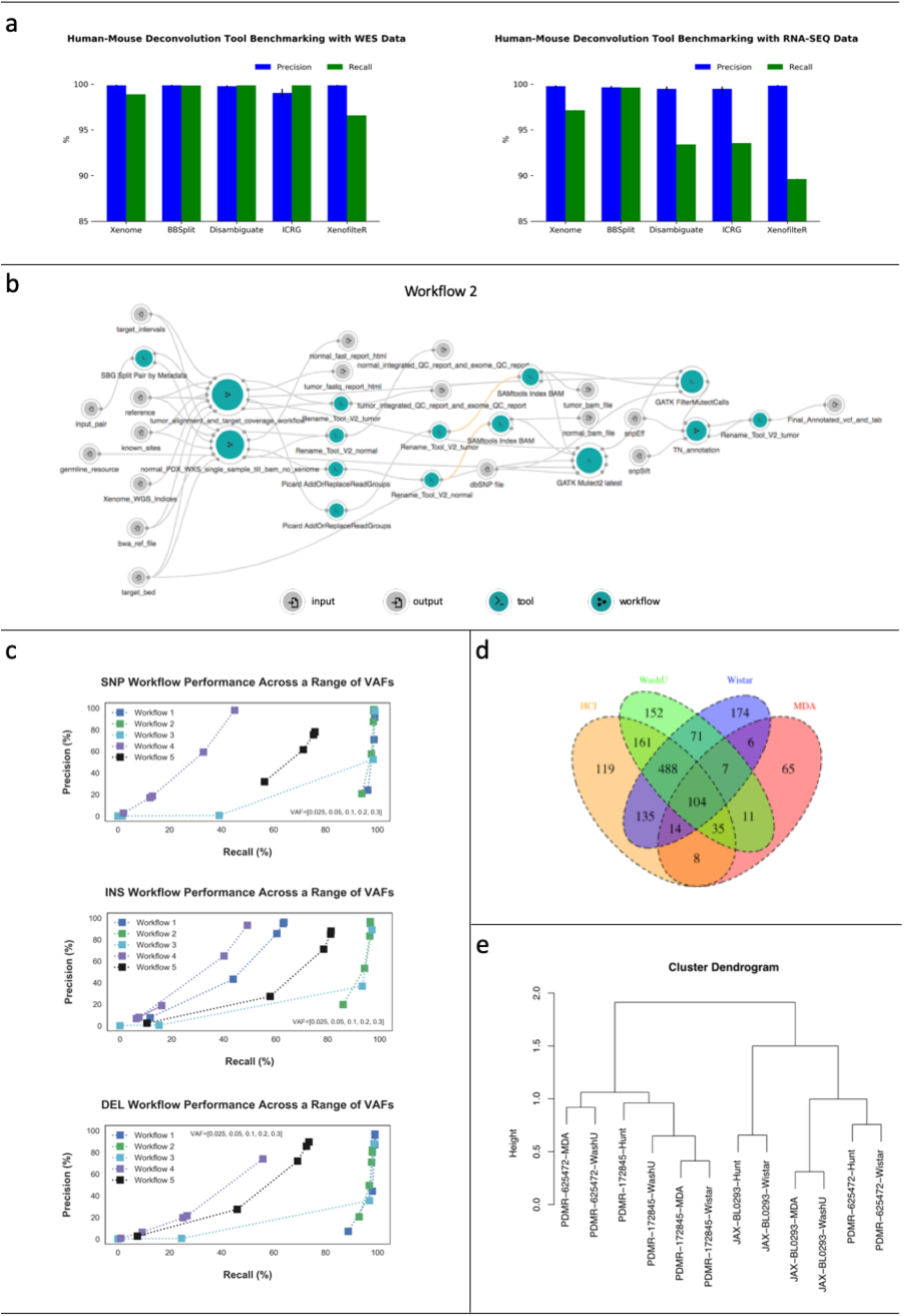
Workflow Benchmarking and Analysis Summary. a) Panel A shows results of the evaluation of mouse-human disambiguation tools (Xenome, BBSplit, Disambiguate, ICRG, XenofilteR). Each figures shows precision (blue) and recall (green) for a simulated data. Left figure shows results of mouse disambiguation for whole exome data. Right figure shows results of mouse disambiguation for RNA-seq data. b) The panel shows the wiring diagram for the whole exome workflow selected to process data for this study. The selected workflow was selected from 5 workflows submitted by the PDX Development Trial Centers. Wiring diagrams for submitted whole exome workflows submitted by the PDX Development and Trials Centers. Wiring diagrams include nodes and connections. Nodes depict inputs -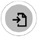, outputs -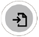, tools - 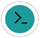, and workflows - 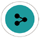. Connections between nodes depict that input to a node is from the output of another node. Orange nodes - 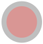 identify a tool or a workflow with an available update. c) Panel shows performance evaluations of five workflows submitted by the PDTC. Each workflow was evaluated by SNP (top), INS (middle), and DEL (bottom) with a range of variant allele frequencies (0.025, 0.05, 0.3, 0.2, 0.3). Each plot shows recall and precision respectively on the x and y axis. Results for each of the workflow are shown with the same color: Workflow 1- blue, Worfklow 2 – green, Workflow 3-light blue, Workflow 4 – purple, and Workflow 5 – black d) A Venn diagram showing the overlap in high-quality variant calls for model JAXBL029 by model using intersected array and removing lower allele frequency (AF) calls. e) Dendrogram of median polish by center (by MBatch) using TMM normalized count per million values

### Benchmarking of WES analysis pipelines

We next compared WES results generated by the five pipelines including variant calling and the effectiveness of mouse-human disambiguation. For this analysis, two simulated benchmark datasets were created, with two levels of mouse contamination (10% and 50%) and a range of variant allele frequencies (VAFs) - 0.025, 0.05, 0.1, 0.2, and 0.3, with spike-ins of point mutations and indels (See Methods). For performance metrics, we used precision/recall (across SNPs, INS, DELs) and pseudo-ROC curves (see Methods). We observed minimal impact of different percentages of mouse contamination on the performance of the five workflows **(Supplementary Table 3)**. When analyzing variant caller performances, we observed that MuTect2 (used in Workflows 2 and 4) performed consistently well across all samples for all the tested VAF levels. **Supplementary Figure 11** shows SNP performance across 0.05 and 0.3 VAFs for BS-DN dataset across different coverage values (although we only show 2 VAF levels, the caller performed well across all VAF levels tested i.e. 0.05 - 0.3); however, indel recall decreased at lower VAFs (Figure 4). VarScan2 (used in Workflows 3 and 4) called only a small number of variants at lower VAFs as evident from the very low recall values depicted in Figure 4. We also observed marked differences in performance of two VarScan2 PDTC workflows, e.g. the DN dataset when processed through workflow 3 at low VAFs i.e. at 0.025 and 0.05 VAF had SNP precision values of 0% and 1.71%, respectively, and when processed through workflow 4 had SNP precision values of 2.16% and 12.4%. The difference in performance between workflows 3 and 4 is possibly due to the fact that in workflow 3 Varscan2 was run independently, whereas in workflow 4 the final calls are a union of VarScan2 and Mutect2 calls. Recall was good at higher VAFs, but precision varied (Figure 4). For example, the DN dataset when processed through workflow 3 at 0.2 and 0.3 VAF had SNP precision values of 98.43% and 99.13%, respectively, and when processed through workflow 4 had SNP precision values of 33.04% and 45.03. Strelka2 (part of workflows 1 and 5) was the most aggressive caller, achieving considerable recall even at the lowest VAFs tested (Figure 4). However, Strelka2 performance varied between the two workflows that used it, i.e. workflow 1 and workflow 5, possibly because workflow 1 used the recommended settings for running Strelka (combining it with Manta), whereas workflow 5 ran Strelka independently. We observed similar trends in the pseudo-ROC curves (not shown) consistent with results described above and depicted in Figure 4 and **Supplementary Figure 11**.

### PDXNet Exome, CNV, and RNA-seq workflows

According to the achieved precision and recall values across SNPs, INS, and DELs (F1 statistic), Workflow 2 was the best performing WES workflow for PDX data. Consequently, we recommend using Workflow 2 for somatic calling in PDX tumor-normal paired WES samples. As the other workflows (**Supplementary Figures 12 and 13**) may be suited for other datasets we are releasing all workflows on the CGC. In addition, we are releasing a tumor-only exome-seq variant calling pipeline, an RNA-seq expression pipeline, and a CNV calling from exome-seq pipeline (**See Methods and Supplementary Figure 14**). The tumor-only exome-seq, RNA-seq, and CNV calling pipelines were used to analyze samples from each PDTC in the temozolomide experiments.

### Robustness of PDX Sequence Evaluations

To test the robustness of these sequence analysis workflows, we applied them to PDX samples from the temozolomide study. Each PDTC generated an independent biological sample of an untreated PDX for each of the three patient models. They then sequenced these independently and submitted the sequence data to the coordinating center.

### Variant Calls from Exome-Seq

FASTQ files from whole exome sequencing were obtained from the four PDTCs (MD Anderson, Huntsman Cancer Institute, Washington University in St. Louis, and the Wistar Institute). Each center provided WES and RNA sequencing data from the PDX models: JAX-BL0293, PDMR-172845 and PDMR-625472. No matched normal data were available for these models.

The WES data were analyzed with the optimal WES pipeline that was modified to take into account the lack of normal DNA, i.e. the ‘PDX WES Tumor-Only: Mutect2’ workflow. The exome capture kits used by each center covered different regions and total amounts of the genome **(Supplementary Table 4)**, resulting in disparate variant calls among centers. (Figure 4). The length of the genome covered by the intersection of the capture loci across all groups was 33.71Mb. Applying these filters made the average number of variant calls across centers for each model comparable (**Supplementary Table 5, Figure 4, Supplementary Figure 15**), though centers with lower sequencing depth had fewer calls meeting the QC threshold. A distribution of allele frequencies for calls meeting the QC threshold for each sample across each center is shown in **Supplementary Figure 16**.

### Copy number calls from Exome-Seq

We called the copy number for each sample using CNVkit with a pooled normal approach (Talevich et al. 2016) (**Supplementary Figure 17**). Overall, we observed similar profiles among samples from the same model. The most apparent difference between samples was an overall shift relative to the baseline. As such, comparing absolute copy number gain and loss calls between samples remains challenging. **Supplementary Figure 18** shows the Pearson correlation coefficients between samples. We observed higher Pearson coefficients (>0.746) for pairwise comparisons for samples of the same tumor among the HCI-BCM, WASHU and Wistar centers, compared to samples of different tumors. On the other hand, the MD Anderson profiles were noisier due to lower coverage, despite us using the “drop low coverage” option in CNVkit, and we were unable to identify strong correlations between samples of the same tumor for the MD Anderson group.

### Expression calls from RNA-Seq

Data provided by each center were generated using different RNA-sequencing protocols (**Supplementary Table 6)** and were analyzed with the rsem-1-2-31-workflow-with-star-aligner (single-end data) and rsem-1-2-31-workflow-with-star-aligner-pe (paired-end data) workflows on the CGC, with small adjustments based on single vs. paired end sequencing or directionality parameters (see Methods). To account for differences in library size, data were normalized by Trimmed Mean of M-values (TMM), and further converted to count per million (CPM) with the R package edgeR ^32,33^. Following normalization and CPM conversion, significant batch effects were still present in these data (**Supplementary Figure 19).** To correct for batch effect among centers, median polish by center was applied to TMM normalized CPM data as implemented in the MBatch R package (github.com/MD-Anderson-Bioinformatics/MBatch). Following batch correction, samples tended to cluster by model rather than sample, though with some exceptions (Figure 4).

## Discussion

Our work demonstrates the robustness of PDXs as a model system for studying cancer drug response. In particular, we have demonstrated the experimental robustness of PDX response for three different models even among research groups blinded to the expected response and who followed independently developed preclinical protocols. This is a key result that shows that, even when groups are not told what experimental protocol to use, PDXs can yield accurate and consistent treatment responses.

In addition, we have developed standardized PDX sequence analysis pipelines for tumor-normal variant calling, tumor-only variant calling, and RNA-seq expression calling. We have provided these as public tools on the CGC, making them easily accessible for other researchers and applicable to the broad data collections shared on the CGC. Not only have these pipelines been tested on extensive benchmark datasets, but we have also applied the tumor-only variant calling and RNA-seq pipelines to sequence data generated across the PDTCs in the temozolomide study. These give similar results across the groups, demonstrating both the efficacy of the pipelines and the minor sequence evolution from PDX to PDX during the process of generating test cohorts across groups.

Importantly, we have also developed biostatistical analysis workflows for tumor volume data, which we are releasing here as well. Our results show a high level of concordance among the various biostatistical analysis strategies, but with some caveats. The RECIST criteria are heavily threshold dependent, has lower statistical power, and less consistent with results from the other strategies. Since each strategy has its own strengths and weaknesses, we recommend testing multiple strategies for PDX analyses. It is also important to consider clinical as well as statistical significance, considering effect sizes to be sure any effect is of sufficient magnitude to be meaningful, a determination that may depend on the clinical context. Classifying PDX volume data into meaningful patient-analogous categories of complete response, stable disease and partial response remains challenging, though this may become possible as datasets with paired clinical and PDX response data increase. In the meantime, our automated analysis scripts, which collate the results and analytical steps into an automated report, provide a standard tool for the PDX field, and future PDXNet volume data will be released in a data format consistent with these scripts. We encourage others to follow the volume data standards we have developed here, which will assist in the quantitative application of PDX treatment data for predicting the efficacy of drugs in patients.

## Methods

### Animal Models

Three PDX models were selected based solely on their temozolomide responsiveness. They were 625472-104-R (colon adenocarcinoma), 172845-121-T (colon adenocarcinoma), and BL0293-F563 (urothelial/bladder cancer). Cryopreserved PDX tumor fragments were shipped from the PDMR to the individual PDTCs, implanted for initial expansion and then passaged for the preclinical study. Briefly, cryopreserved PDX material was prepared into implantation size pieces as outlined in Table 1. The PDX material plus a drop of Matrigel (BD BioSciences, Bedford, MA.) was then implanted subcutaneously in NOD.Cg-Prkdc^scid^ Il2rg^tm1Wjl^/SzJ (NSG) host mice. Mice were housed in sterile, filter-capped polycarbonate cages, maintained in a barrier facility on a 12-hour light/dark cycle, and were provided sterilized food and water, ad libitum. Animals were monitored weekly for tumor growth. The initial passage of material was grown to approximately 1000-2000mm^3^ calculated using the following formula: weight (mg) =(tumor length x [tumor width]2)/2 ^34^. Tumor material was then harvested, a portion cryopreserved, and the remainder implanted into NSG host mice for the preclinical drug study. Related patient data, clinical history, representative histology and short-tandem repeat profiles for the PDX models can be found at https://pdmr.cancer.gov; model BL0293-F563 was originally developed by The Jackson Laboratory (tumor model TM00016, http://tumor.informatics.jax.org/mtbwi/pdxSearch.do).

### Preclinical Studies

Specific tumor staging size, implantation method, and cohort size at the PDMR and each PDTC are outlined in Table 1 based on each site’s standard practices. In general, tumors were staged to a preselected size (weight = 100-200 mg). Tumor-bearing mice were randomized before initiation of treatment and assigned to each group. Body weight was monitored 1-2 times weekly and tumor size was assessed 2-3 times weekly by caliper measurement. For all sites, drug studies were performed at passage 3 for 625472-104-R, passage 4 for 172845-121-T, and passage 6 for BL0293-F563 (passage 0 = first implanted host). Temozolomide (NSC 362856) was obtained from the Developmental Therapeutics Program, NCI and administered at the times and doses indicated in Table 1. Animals were sacrificed when the tumors reached an individual PDTC’s animal welfare endpoint or a maximum tumor size; if tumor growth delay was observed a tertiary endpoint was used by some sites (Table 1).

### Ethics Statement

The Frederick National Laboratory for Cancer Research (location of the PDMR) is accredited by the Association for Assessment and Accreditation of Laboratory Animal Care International and follows the USPHS Policy for the Care and Use of Laboratory Animals. All the studies were conducted according to an approved animal care and use committee protocol in accordance with the procedures outlined in the “Guide for Care and Use of Laboratory Animals” (National Research Council; 1996; National Academy Press; Washington, D.C.).

All patients and healthy donors gave written informed consent for study inclusion and were enrolled on institutional review board-approved protocols of record for the sites that developed the PDX models (DCTD, NCI and The Jackson Laboratory). The study was performed in accordance with the precepts established by the Helsinki Declaration. The study design and conduct complied with all applicable regulations, guidances, and local policies and was approved by the institutional review board of record for each PDTC

### Computational Workflows

All analyses were performed on the CGC (https://cgc.sbgenomics.com/) ^35^ with workflows and tools implemented using Common Workflow Language. Human and mouse data were aligned to GRCh38 and mm10 assemblies, respectively. All workflows are available in the Temozolomide Pilot Workflows Project on the CGC. CGC users can request access to the workflows by emailing.

#### Human-mouse read deconvolution

For the WES data benchmark, an experimental WES series of simulated human-mouse mixtures (90:10, 80:20, 60:40, 50:50, 40:60, 20:80) was created by mixing two 100 % pure WES human and mouse samples (sample ids 14311X2 and 14311X8, respectively). A Python script based on HTseq (v0.6.1) was used to create the mixtures by randomly subsampling input pairs of human and mouse FASTQ.GZ files to a predefined fraction to obtain 56-57 million read pairs per data point. We also tested RNA-seq data using a simulated human-mouse mixture series (90:10, 80:20, 60:40, 50:50, 40:60, 20:80) based on a pair of human and mouse lung tissue RNA-seq samples (ENCSR129KCJ and ENCSR870AQU, respectively) ^36^. The RNA-seq datasets were prepared with the same Python script used for mixing WES data.

We compared several tools for mouse-human read deconvolution. These were Xenome (v1.0.0) ^37^, BBSplit (v37.93) ^38^, Disambiguate (v1.0; commit c52402a) ^38^, ICRG (Callari et al., 2018) ^39^, and XenofilteR (v1.5) ^40^. BBtools Seal tool was also initially considered; however, it was excluded because of extremely high default RAM requirements. These methods followed three main approaches: BBSplit and Xenome operate on raw FASTQ data, with BBSplit aligning whole reads and Xenome using k-mers to classify reads. Disambiguate and XenofilteR require pre-aligned inputs (i.e. reads aligned to both host and graft reference genomes) and use alignment quality scores for classification. ICRG relies on aligning reads to a combined host-graft reference sequence.

For tools requiring aligned data inputs (BAM files), BWA-MEM ^40^ was used for alignment. For the mouse-human disambiguation step, only reads unambiguously classified as human by a tool were labeled “human.” All other reads were considered “not human” for the true/false positive/negative calling. This classification scheme provided a simplified common framework for evaluating all benchmarked tools. All metrics were calculated via a Python script based on HTseq.

#### Tumor-normal WES variant calling

##### Benchmark Datasets

Two simulated whole exome-seq datasets were used in the benchmark for the tumor-normal variant calling workflow. The first dataset (DN) was prepared by researchers from Huntsman Cancer Institute and consisted of 100x data based on two normal samples, spiked with 30,466 SNPs, 1,723 insertions, and 4,192 deletions sampled from ClinVar, at 0.025, 0.05, 0.1, 0.2, and 0.3 simulated variant allele frequency (VAF) and 10 and 50 % mouse contamination. Germline variants were called with HaplotypeCaller and blacklisted before analysis. The second dataset (BS) was NA12878 WES data (~250x coverage; with 10 % mouse reads contamination) which was spiked with BamSurgeon [i] (default parameters, haplosize 151) at 0.05, 0.1, 0.2, and 0.3 VAF using both the ClinVar variant set used for the other simulated dataset (BS-DN) and 30000 TCGA BRCA SNPs combined with indels from the ClinVar set) (BS-BRCA). For both variant sets, 0.05 and 0.3 VAF samples were also downsampled to 130x and 65x to analyze coverage effects (the experimental coverage of the datasets was 250x).

##### Workflow testing

Five tumor-normal WES data analysis workflows from PDXNet research groups were tested on the benchmark sets, as detailed in **Supplementary Table 2**, with the goal of evaluating the accuracy in the presence of variable mouse contamination, coverage, and VAF. Starting from FASTQ data the workflows performed mouse-human disambiguation, alignment, and variant calling with one or more somatic variant callers. For the variant calling step, Mutect2 ^41,42^, VarScan2 ^43^, and Strelka2 ^14^ each featured in two workflows. Manta ^44^ and Pindel ^45^ structural variant callers were also used, but were not evaluated, as the benchmark focused on small variants, i.e. SNPs and indels with length <50 base pairs.

Precision/recall and pseudo-ROC curves were used to evaluate the detection of SNPs, insertions and deletions. Pseudo-ROC curves are plots of descending false discovery rate (FDR) vs. true positive rate (recall). To produce the curves, VCF calls are ranked using a caller-provided quantitative score. These were TLOD, SSC, QSS/QSI for MuTect2, VarScan2 and Strelka2, respectively. These rankings allowed us to order the calls for the FDR and TPR calculations. Python scripts were used to calculate the relevant metrics.

For all the submitted workflows, default parameters were used as specified by the workflow authors. Details are provided in the workflows that are accessible through the CGC upon request.

#### Tumor-only WES variant and CNV calling

Because a substantial number of PDXs among the broader research community lack matched normal DNA, we also developed a workflow for tumor-only mutation calling **(Supplementary Figure 14)**. All reads were required to meet the quality control cutoff that at least half of the nt positions have >20 phred base quality. We removed adaptors using Trimmomatic v 0.36 (Bolger et al. 2014). Trimmed reads were aligned to the human genome (build GRCh38.p5) using bwakit v0.7.15 (Li and Durbin 2009). Mouse reads were removed with xenome v 1.0.0 (Conway et al. 2012) at default parameters. The alignments were converted to BAM format using Picard SortSam v 1.140 (https://broadinstitute.github.io/picard/), and duplicates were removed by the Picard MarkDuplicates utility. BaseRecalibrator from the Genome Analysis Tool Kit (GATK) v4.0.5.1 ^46,47^ was used to adjust the quality of raw reads. Training files for the base quality scale recalibration were Mills_and_1000G_gold_standard.indels.hg38.vcf.gz, Homo_sapiens_assembly38.known_indels.vcf.gz, and dbSNP v151. Mean target coverage was determined for each sample by Picard CollectHsMetrics, and a MultiQC ^48^ report was generated. Aligned BAM files were indexed by GATK and passed to GATK Mutect2 v 4.0.5.1. Variants were called in Mutect2 using the Exome Aggregation Consortium ^49^ database lifted over to GRCh38 as a germline reference with the allele frequency of samples not in reference set to 0.0000082364. Variant calls were then filtered using GATK FilterMutectCalls v 4.0.5.1. Calls were segregated by chromosome with SnpSift Split v 4.3 ^50^ and annotated by SnpEff v 4.3 ^51^ using the snpEff v4.3 GRCh38 database. Two additional annotations of variant calls were done with SnpSift dbNSFP using database dbNSFP3.2a, and with SnpSift Annotate using the catalogue of somatic mutations in cancer (COSMIC) v80 database ^52^. A reference implementation of this workflow is developed and deployed on the CGC.

To call copy number, we built a pooled normal reference using CNVkit v0.9.3 ^53^ from the three samples that used the same exome-seq capture kit and with sex matching. Afterward we used CNVkit to call the CNV segments from each sample using the pooled normal reference. The MD Anderson samples exhibited low mean target coverage so we turned on the --drop-low-coverage option in CNVkit to reduce the noise in the CNV profile.

#### RNA-seq expression calling

Because the disambiguation of mouse and human reads was sharp for both DNA and RNA data, we did not expect expression calling workflows to have issues specific to PDXs. Therefore, we dockerized only one PDX RNA-seq expression workflow **(Supplementary Figure 14)** that was submitted by The Jackson Laboratory (JAX). The transcriptomes of hg38 and NOD (based on the mm10 mouse genome) were used to construct the xenome (version 1.0.0) ^37^ indices (k=25), and then reads were classified as human, mouse, both, neither or ambiguous at default xenome parameters. Reference indices for the alignment were built by rsem-prepare-reference using ENSEMBL annotation (version GRCh38.91) for STAR aligner (version 2.5.1b) ^54^. Human-specific reads were mapped to reference indices using STAR, and expression estimates were computed using rsem-calculate-expression v1.2.31 ^33^ at default parameters. Picard CollectRnaSeqMetrics: (broadinstitute.github.io/picard/picard-metric-definitions.html) was used to calculate the post-alignment mapping statistics. An implementation of this workflow has been deployed on the CGC.

#### Comparisons of xenograft sequence data across PDTCs

Each PDTC submitted WES and RNA-seq data from untreated xenografts that had been successfully grown in mice at the respective sites. Groups were asked to submit xenograft sequence data according to their standard practices, without pre-specification of the sample passage number or the sequencing protocol. In the intersection analysis, only variants with allele frequency > 0.2 were retained. We note that MD Anderson had fewer calls that passed the allele frequency filter in comparison to other centers. This is because MD Anderson provided samples had mean target coverage ~30X whereas samples from other centers were sequenced to a depth of ~150X (**Supplementary Table 7**).

For the copy number comparisons, the copy number alteration (CNA) segments obtained from CNVkit using a pooled normal were median-centered and visualized in IGV v2.4.13 ^55^. To determine the overall concordance of the CNA between each pair of samples, we first intersected the CNA segments for each pair of samples and then binned them into 100kb-windows using Bedtools v2.26.0 ^56^.

RNA-seq data provided by each center were generated using different kits and protocols, and the data from Huntsman institute was sequenced in single end mode **(Supplementary Table 6)**. Sequence data were analyzed with the ‘PDXnet RNA Expression Estimation’ and the ‘PDXnet RNA Expression Estimation – SE’ workflows on the CGC. RNA expression estimates were downloaded from CGC for additional analyses. The single-end data provided by Huntsman yielded estimates of RNA expression that were twice as high when compared to the paired-ended sample provided by other centers due to differential handling of paired-end and single-end data by RSEM ^33^ tool. To eliminate the biases in the count estimation across centers, Huntsman, estimated transcript counts were divided in half. From the mapping stats and from automatic library type detection algorithm in the tool Salmon, we noted that RNA-Seq library generated at MDAnderson are non-directional though the sequencing protocol used is for directional library thus we decided to consider MDAnderson library as non-directional during the analysis.

### Statistical Analysis of Tumor Growth Data

There is not a single consensus in literature in terms of which endpoint to use to measure tumor response in PDX models. There are a number of potential options. Rather than considering just one, our strategy was to consider a wide range of potential analytical strategies, each of which captures different aspects of the response and has its own strengths and weaknesses. Here, we compare and contrast them in this pilot study and assess the robustness of sensitivity assessments across different analytical strategies, with the goal of making recommendations for the broader community. Towards this goal, we built an R analysis pipeline that computes all of the following measures as well as generates a set of useful graphical summaries.

#### Percent Change in tumor volume 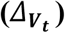

For an individual mouse, the response was determined by computing percent change in tumor volume from baseline to time as follows: % tumor 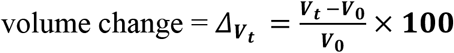, where *V*_*t*_ is the tumor volume at time t and *V*_*0*_ is the tumor volume at baseline. For animals for which there is no tumor volume measurement at time t but which have flanking volume measurements at time *t*_0_ and *t*_1_ such that *t*_0_ < *t* < *t*_1_, then we use linear interpolation to compute the measurement. That is, we compute

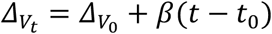

Where 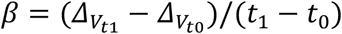.

All responses defined below were based on the interpolated tumor volume changes.

#### Area under the tumor growth curve up to time t (α*AUC*_*t*_)

For this measure, we computed the area under the tumor growth curve from baseline up to time t, normalized by dividing by t. With this normalization factor, the interpretation of this measure is the average percent change in tumor size from baseline to time. If there was no tumor measurement at time t but measurements at flanking measurements at time *t*_0_ and *t*_1_ such that *t*_0_ < *t* < *t*_1_, then we used linear interpolation to estimate the tumor volume at t.

#### Adjusted area under the curve (α*AUC*_*max*_)

We also computed the area under the curve from baseline to the last measurement time point, which for animal i is given by notation τ_*i*_. We adjusted the AUC by dividing by the length of the interval between the baseline and the last time point for which a tumor volume was computed for each mouse,τ_*i*_, which makes the interpretation of this measure the average percent change in tumor size from baseline to last measurement in the study.

#### RECIST criteria (*RECIST*_*t,c*_)

This outcome mimics the typical RECIST response criteria commonly used in clinical trials for solid tumors to classify each patient as complete response (CR), partial response (PR), stable disease (SD), or progressive disease (PD). At prespecified time t, each animal was classified into one of these four categories based on their percent tumor volume change from baseline based on cut points *c*_1_ < *c*_2_ < *c*_3_, with complete response (CR) when 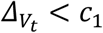; partial response (PR) when 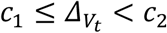, stable disease (SD) when 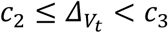, and progressive disease (PD) when 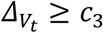.

We considered various sets of cut points (*c*_1_, *c*_2_, *c*_3_), including (−95,−50,10), (−95,−30,20), (−95,−30,50), (−95,−30,100), (−95,−50,50) and (−95,−50,100), and for our analyses in this paper we computed these at time t=21 days.

#### Tumor Growth Inhibition (*TGI*_*t*_)

To measure antitumor activity of the treatment group compared to the control group, we considered the tumor growth treatment-to-control ratio (gamma_t) estimated by one way ANOVA focused on time t. Let *R*_*it*_ be, for animal i, the ratio of tumor size from baseline to time t, i.e. *R*_*it*_ = *V*_*i0*_/*V*_*it*_ where *V*_*i0*_ and *V*_*it*_are the tumor volumes for animal i at baseline and time t, respectively. Let *T*_*i*_ be an indicator of whether animal i was given the active treatment (*T*_*i*_ =1) or control (*T*_*i*_ =0). After log transforming the ratios, we fit the following linear model:

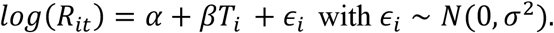

If we define 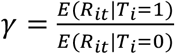 as the ratio of mean tumor/mean control, called the tumor growth inhibition it can be shown that

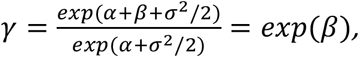

The degree to which this measure is less than 1 indicates the degree of growth inhibition of the treatment. Thus, to test for antitumor activity of the treatment group at time t, we can test the null hypothesis of no treatment effect by comparing

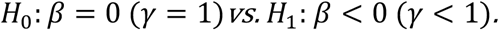

#### Time-to-Event Survival Analysis

Many different events can be evaluated with this method. For example “time to tumor doubling/tripling/quadrupling”, “time to tumor halving”, “time to complete response” etc. depending on the growth characteristics of the PDX collection in question. In this study, the event was defined as the time until the tumor volume increases by a multiple of *δ*, 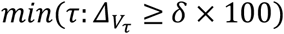(e.g. *δ* = 4 corresponds to time until tumor quadruples in size). This measure will be censored at the last measurement value for animals whose tumors never increased by that multiple.

#### Statistical analysis

One-way ANOVA or two-sample t-tests were performed to test the difference of tumor Volume changes 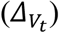 at day t=21 between treatment and control groups as appropriate, and similar analyses were done for the AUC measures. Fisher’s exact test was performed to test the association between treatment and drug response (non-PD vs. PD). The log-rank test was used to compare PFS distributions between treatment and control groups. All of the analysis was implemented using R.

We have developed an R markdown script that can be used to automatically run these analyses and produce summary plots given the input data is formatted as described in the Supplementary Materials. to request the R script that we freely share with this publication for other researchers to use to analyze their PDX data.

## Acknowledgements

Support for this work included funding provided by the NIH to the PDXNet Data Commons and Coordination Center (NCI U24-CA224067), to the PDX Development and Trial Centers (NCI U54-CA224083, NCI U54-CA224070, NCI U54-CA224065, NCI U54-CA224076, NCI U54-CA233223, and NCI U54-CA233306) and to the National Cancer Institute Cancer Genomics Cloud (HHSN261201400008C and HHSN261201500003I).

## Author Contributions

**Analysis**:

AS, DAD, IS, JR, MJH, ML, NM, RP, SA, XW, YAE, ZZ

**Analysis Design**:

AS, CF, DAD, JHC, JR, JR, JSM, LC, MJH, PNR, YAE

**Conducted Experiments**:

AK, ALW, BEW, HC, JW, MD, MX, SD, VR

**Data Collection and Preparation**:

AS, RJ, SN, TS

**Experimental Design**:

AK, ALW, BAV, BEW, BF, DN, FM, JR, JW, LD, MD, MH, MTL, MX, RG, SD, SL, VR

**Interpretation**:

AS, DADI, IS, JHC, JSM, YAE

**Study Design**:

BD, CB, JAM, JD, JHC, JHD, JSM, PNR, YAE

**Workflow Development**:

JR, ML, NM, XW

**Writing Team**:

AS, DAD, IS, JHC, YAE

**Writing Team**:

AS, DAD, IS, JHC, YAE

