## Supplements for "Systematic Establishment of Robustness and Standards in Patient-Derived Xenograft Experiments and Analysis"

### Supplementary information

### Supplementary Materials 1: Tumor volume data Analysis using R

Our analysis scripts for the tumor volume data assume data with a specific format, as outlined here. Given data in this format, our scripts will automatically generate the analyses presented in the methods section as well as certain plots, as outlined below.

#### Input Data

1) Model name
2) Treatment
   : "Control" indicates the control group
(3) Mice ID
(4) Date
   : "mm/dd/yyyy" format
   : The earliest data per animal is the treatment start date, and we assume the tumor volume value is available at this date
(5) On Treatment
   : include two categories (Yes or No)
   : "Yes" means "receiving treatment" including treatment start
(6) Tumor Volume
   : Baseline values should be included.
(7) Body Weight
   : Plan for Toxicity analysis if applicable

Note:

- The column names need to match those in the following example table
- **Baseline tumor volume** value is essential per animal.
- For the statistical tests, each treatment group should at least three samples, and the last Treatment Date at least extend to 21 days (or whatever timepoint is chosen) later than the initial Treatment Date per mouse
- Although we perform the analysis for each PDX model, multiple models can be included into the same data file (stacked by row)
- Optional: for displaying the treatment schedule with labeling of the drug names, drug table needs to be provided

**Supplementary Table 1**. Example data in the specific input format for tumor volume data analysis.

| **Model** | **Treatment** | **Mice ID** | **Date** | **On Treatment** | **Tumor Volume** | **Body Weight** |
|---|---|---|---|---|---|---|
| TC250 | Control | 0 | 9/15/2017 | | 219.375 | |
| TC250 | Control | 0 | 9/18/2017 | | 161.84 | |
| TC250 | Control | 0 | 9/20/2017 | | 516.096 | |
| TC250 | Control | 0 | 9/25/2017 | | 831.744 | |
| TC250 | Control | 0 | 9/28/2017 | | 1062.5 | |
| TC250 | Control | 0 | 10/2/2017 | | 1653.752 | |
| TC250 | Control | 0 | 10/5/2017 | | 1730.416 | |
| TC250 | Control | 0 | 10/9/2017 | | 1849.848 | |
| TC250 | Control | 0 | 10/12/2017 | | 1885.963 | |
| TC250 | AMG232 + AZD6244 | 4406 | 10/10/2017 | | 118.354 | |
| TC250 | AMG232 + AZD6244 | 4406 | 10/12/2017 | | 159.3595 | |
| TC250 | AMG232 + AZD6244 | 4406 | 10/16/2017 | | 314.928 | |
| TC250 | AMG232 + AZD6244 | 4406 | 10/19/2017 | | 155.682 | |
| TC250 | AMG232 + AZD6244 | 4406 | 10/23/2017 | | 47.068 | |
| TC250 | AMG232 + AZD6244 | 480 | 10/10/2017 | | 112.694 | |

#### Instructions for running tumor volume analysis R scripts

A user can execute the tumor volume analysis script within R studio after setting the project title. project path, data file, and the number of days to include in the analysis. The tumor volume analysis script is available from the CGC ( Email cgc@ sbgenomics.com to request access).

### Supplementary Materials 2: Comparison of Growth kinetics of PDX models at each site

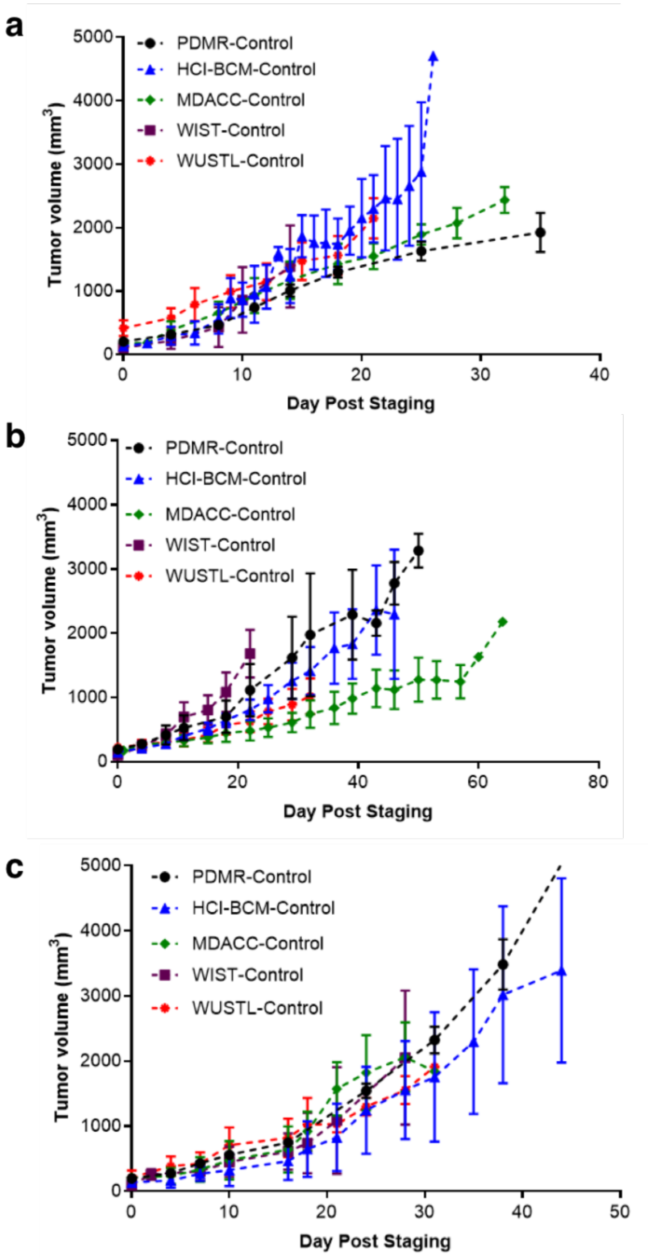

**Supplementary Figure 1**: Vehicle control arms at the PDMR and each PDTC for PDX models 625472-104-R **a)**, 172845-121-T **b)**, and BL0293-F563 **c)**. Data represent median tumor volume ± SD from staging of the drug study (day 0) until end-of-study. Tumor volume is not normalized.

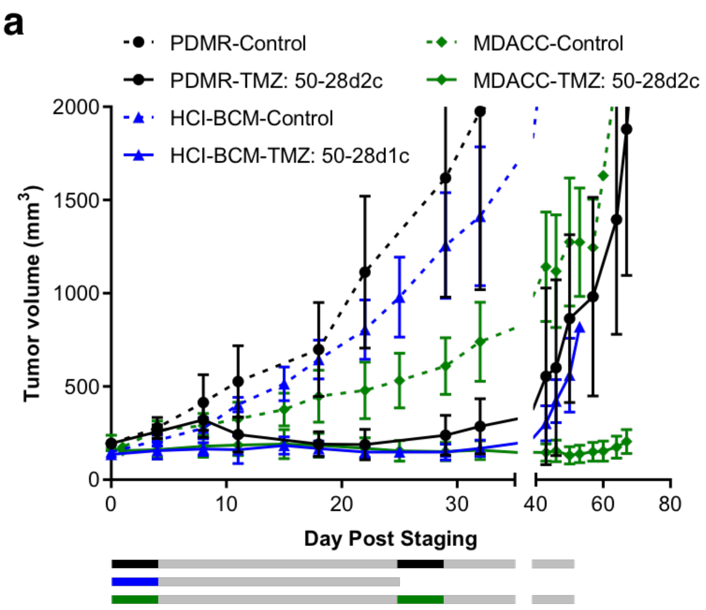

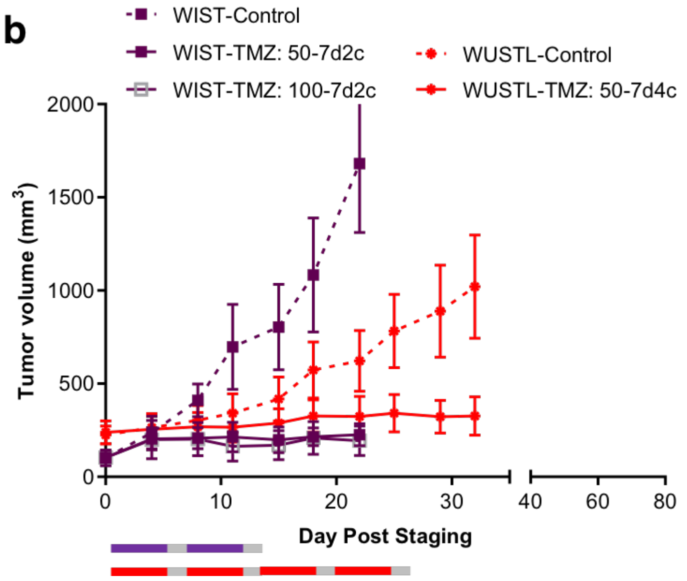

**Supplementary Figure 2:** Comparison of 28-day and 7-day cycle treatment in the intermediary responsive model 172845-121-T. Comparison of tumor volumes observed in **a)** cohorts treated with temozolomide daily for 5 days on a 28-day cycle (1 or 2 cycles) compared to **b)** cohorts treated with temozolomide daily for 5 days on a 7-day cycle (2 or 4 cycles). Colored bars represent treatment cycles at each site. All studies achieved tumor growth inhibition irrespective of cycle length. Dashed lines, vehicle control groups, Solid lines, temozolomide treatment groups. Median ± SD.

**Supplementary Materials 3: Drug response assessment of each PDX model at different sites**

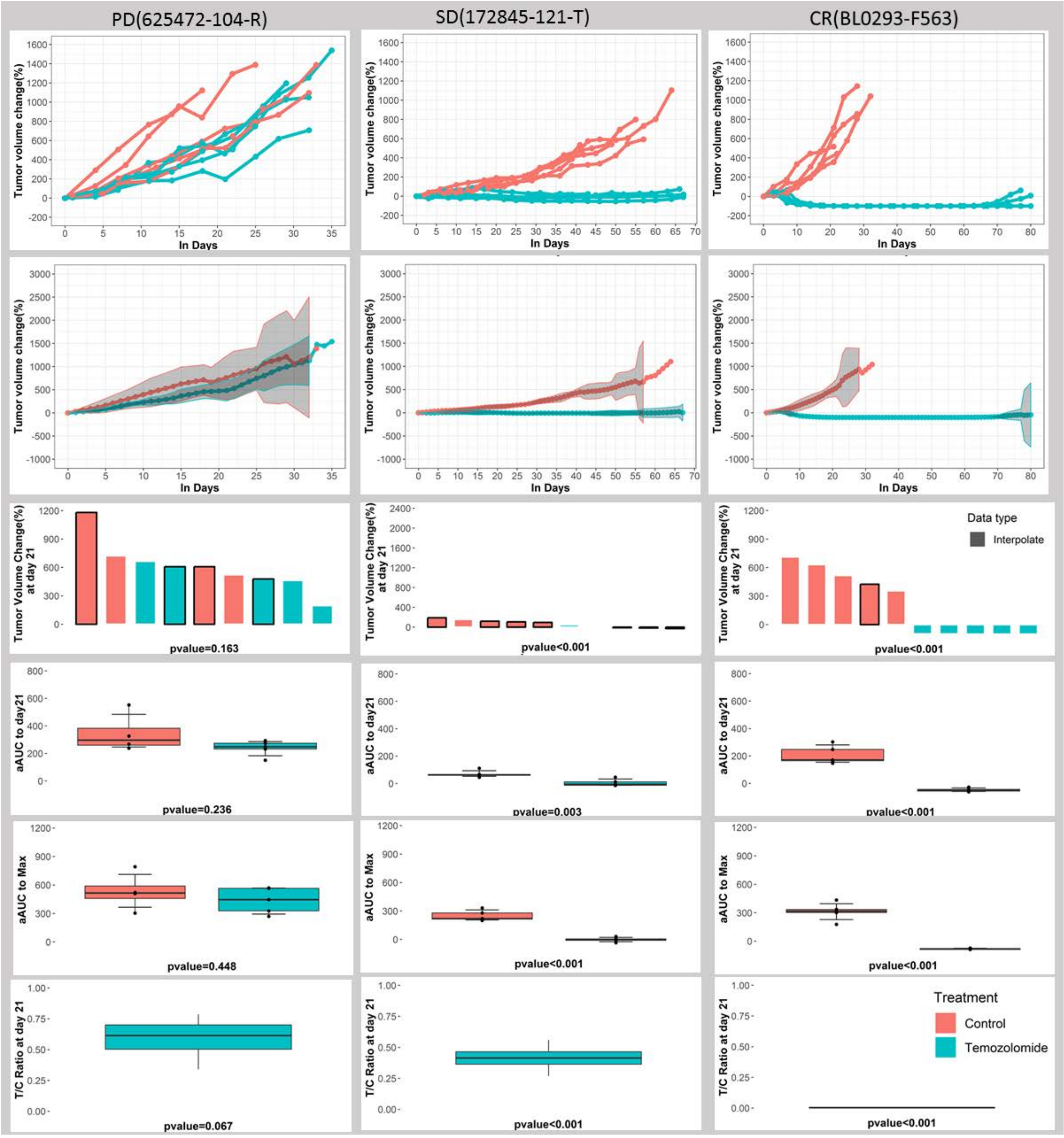

**Supplementary Figure 3.** Analytical Summaries, MD Anderson Study. Analytical results from the MD Anderson study for progressive model (625472-104-R), stable disease model (172845-121-T) and complete response model (BL0293-F563) (columns 1-3, respectively), with interpolated individual curves (row 1), mean curves for treatment and control with 95% confidence bands (row 2), waterfall plots demonstrating $\boldsymbol{\Delta V_{21}}$(row 3), boxplots of $\boldsymbol{aAUC_{21}}$(row 4) and $\boldsymbol{aAUC_{max}}$(row 5) for treatment and control, and a boxplot of $\boldsymbol{TGI_{21}}$(row 6), along with p-values comparing treatment to control for each measure.

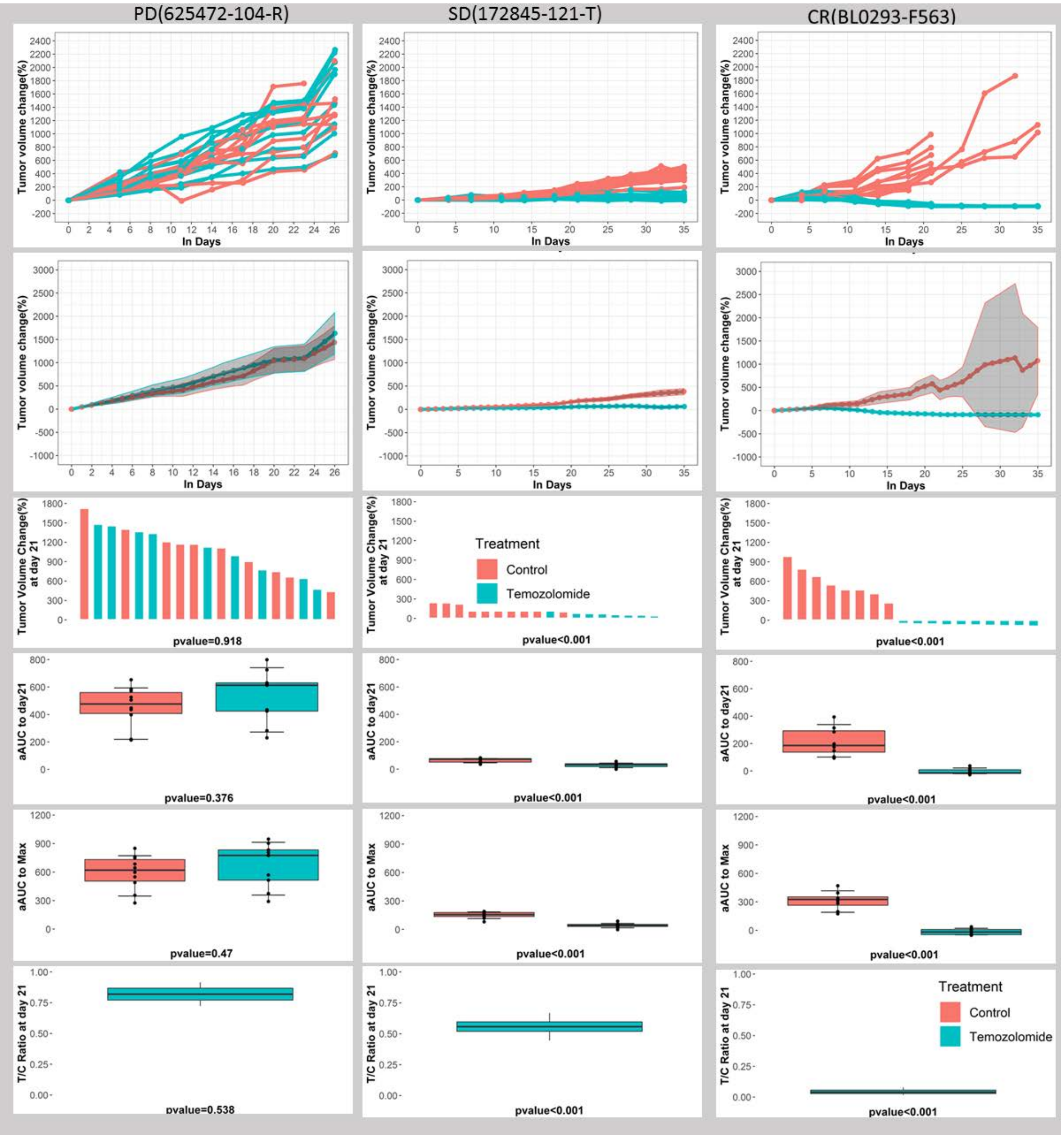

**Supplementary Figure 4.** Analytical Summaries, Washington University Study. Analytical results from the Washington University study for progressive model (625472-104-R), stable disease model (172845-121-T) and complete response model (BL0293-F563) (columns 1-3, respectively), with interpolated individual curves (row 1), mean curves for treatment and control with 95% confidence bands (row 2), waterfall plots demonstrating $\boldsymbol{\Delta V_{21}}$(row 3), boxplots of $\boldsymbol{aAUC_{21}}$(row 4) and $\boldsymbol{aAUC_{max}}$(row 5) for treatment and control, and a boxplot of $\boldsymbol{TGI_{21}}$(row 6), along with p-values comparing treatment to control for each measure.

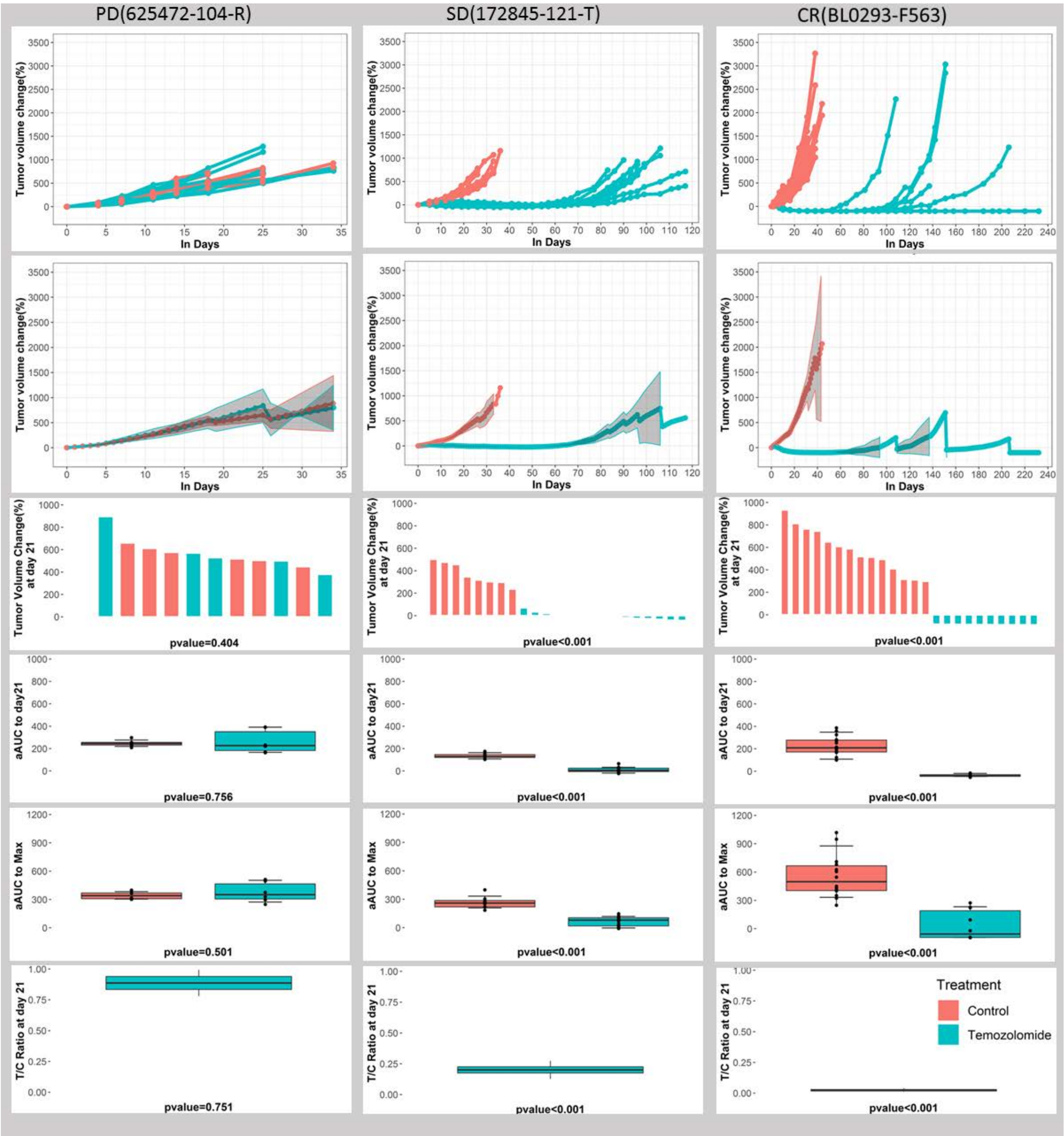

**Supplementary Figure 5.** Analytical Summaries, PDMR Study. Analytical results from the PDMR study for progressive model (625472-104-R), stable disease model (172845-121-T) and complete response model (BL0293-F563) (columns 1-3, respectively), with interpolated individual curves (row 1), mean curves for treatment and control with 95% confidence bands (row 2), waterfall plots demonstrating $\boldsymbol{\Delta V_{21}}$(row 3), boxplots of $\boldsymbol{aAUC_{21}}$(row 4) and $\boldsymbol{aAUC_{max}}$(row 5) for treatment and control, and a boxplot of $\boldsymbol{TGI_{21}}$(row 6), along with p-values comparing treatment to control for each measure.

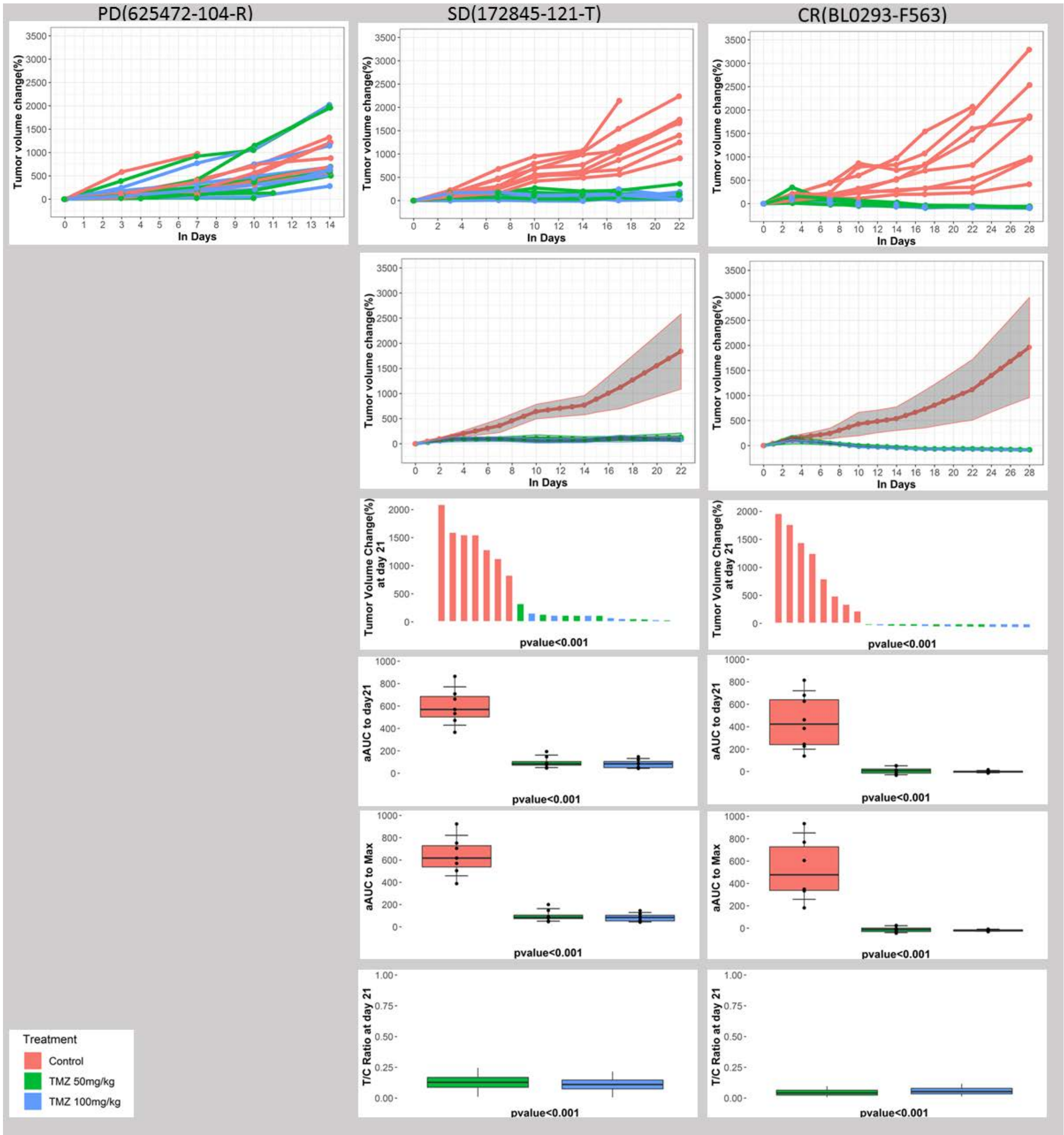

**Supplementary Figure 6.** Analytical Summaries, Wistar Study. Analytical results from the Wistar study for progressive model (625472-104-R), stable disease model (172845-121-T) and complete response model (BL0293-F563) (columns 1-3, respectively), with interpolated individual curves (row 1), mean curves for treatment and control with 95% confidence bands (row 2), waterfall plots demonstrating $\boldsymbol{\Delta V_{21}}$(row 3), boxplots of $\boldsymbol{aAUC_{21}}$(row 4) and $\boldsymbol{aAUC_{max}}$(row 5) for treatment and control, and a boxplot of $\boldsymbol{TGI_{21}}$(row 6), along with p-values comparing treatment to control for each measure.

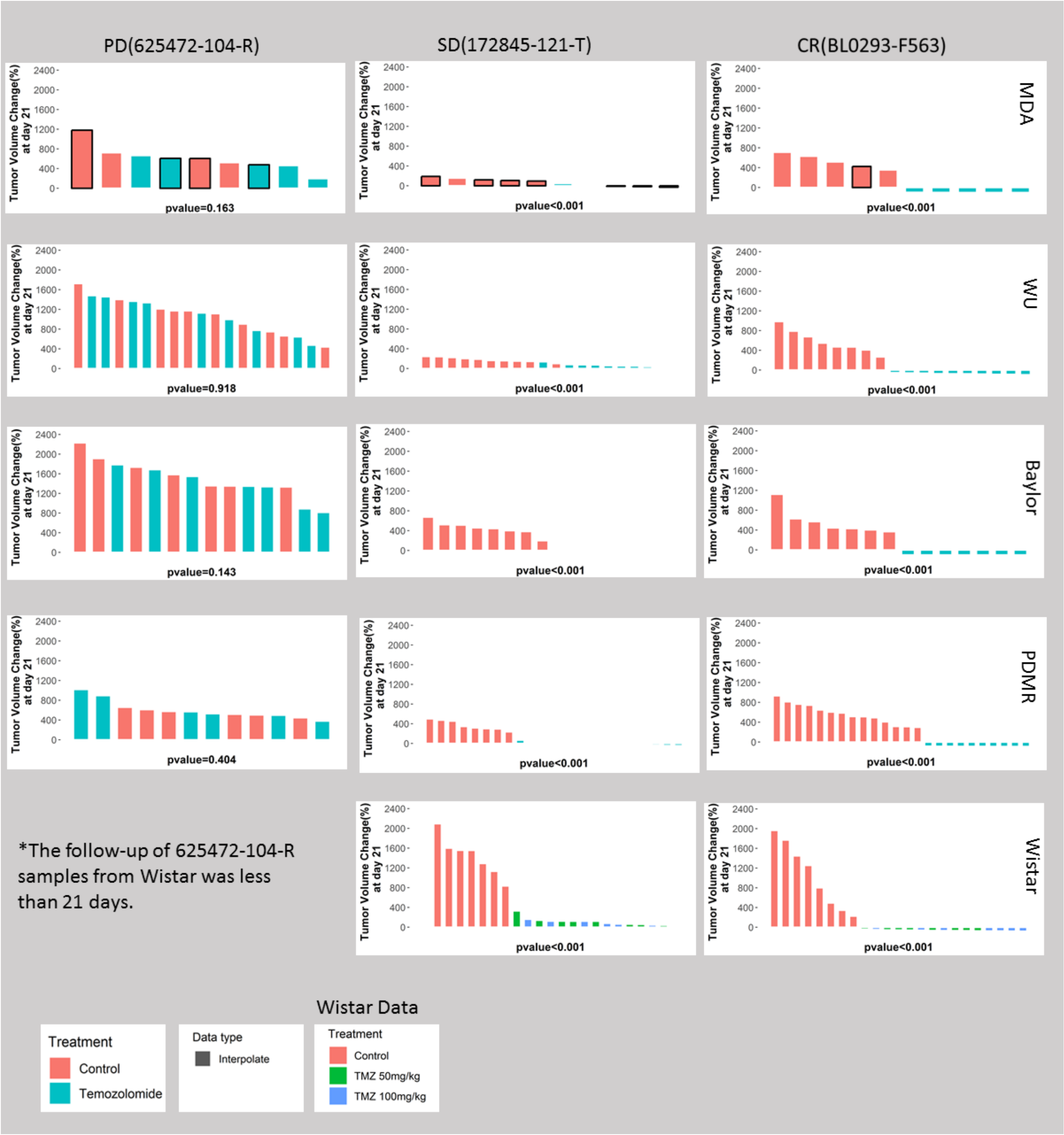

**Supplementary Figure 7.** Results, Analysis of Change in Tumor Volume $\boldsymbol{\Delta V_{21}}$. Analytical results of the change in tumor volume from baseline to day 21 from various studies for progressive model (625472-104-R), stable disease model (172845-121-T) and complete response model (BL0293-F563) (columns 1-3, respectively).

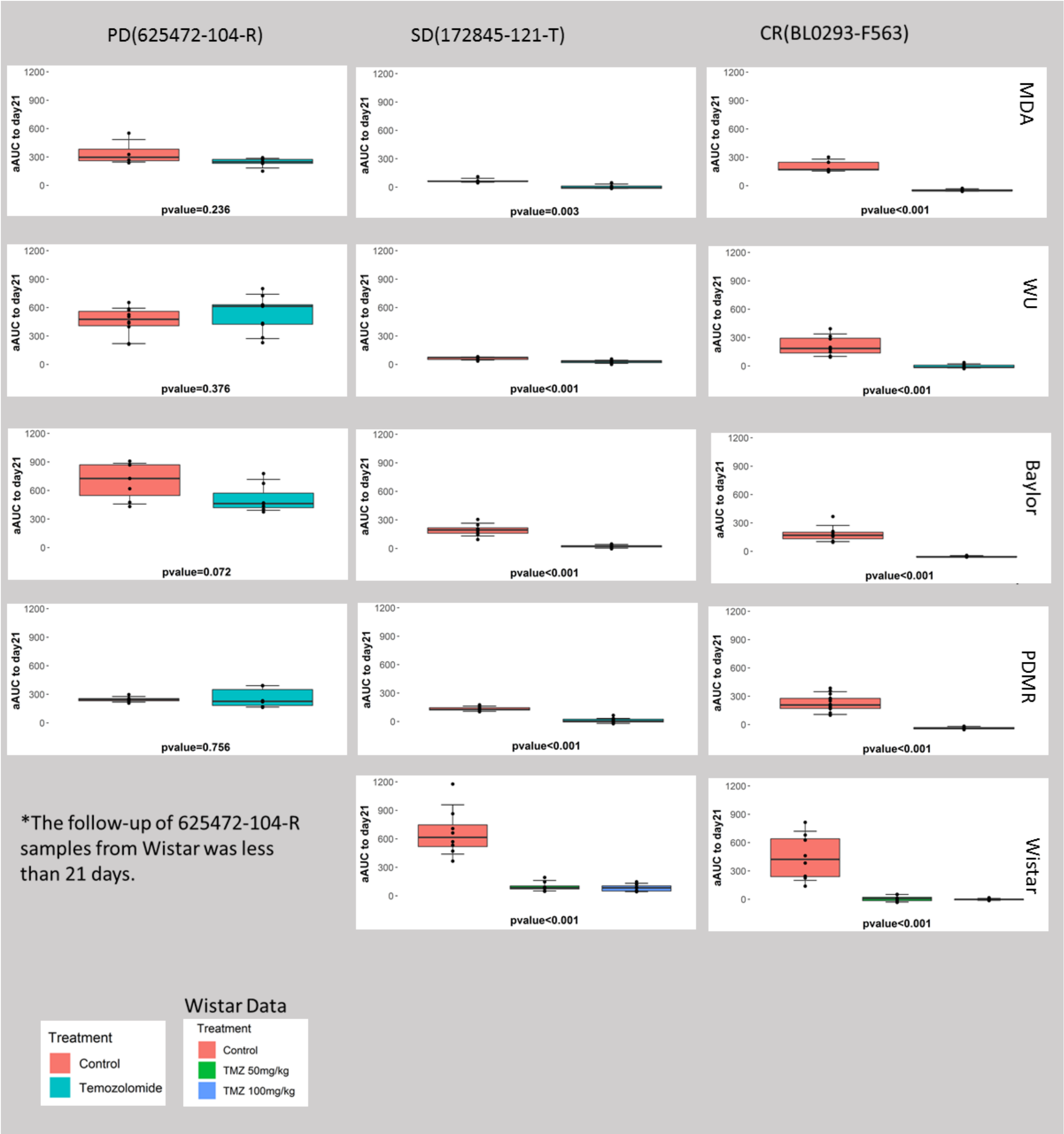

**Supplementary Figure 8.** Results, adjusted area under the curve to 21 days $aAUC_{21}$. Analytical results of the adjusted area under the tumor growth curve from baseline to day 21 from various studies for progressive model (625472-104-R), stable disease model (172845-121-T) and complete response model (BL0293-F563) (columns 1-3, respectively).

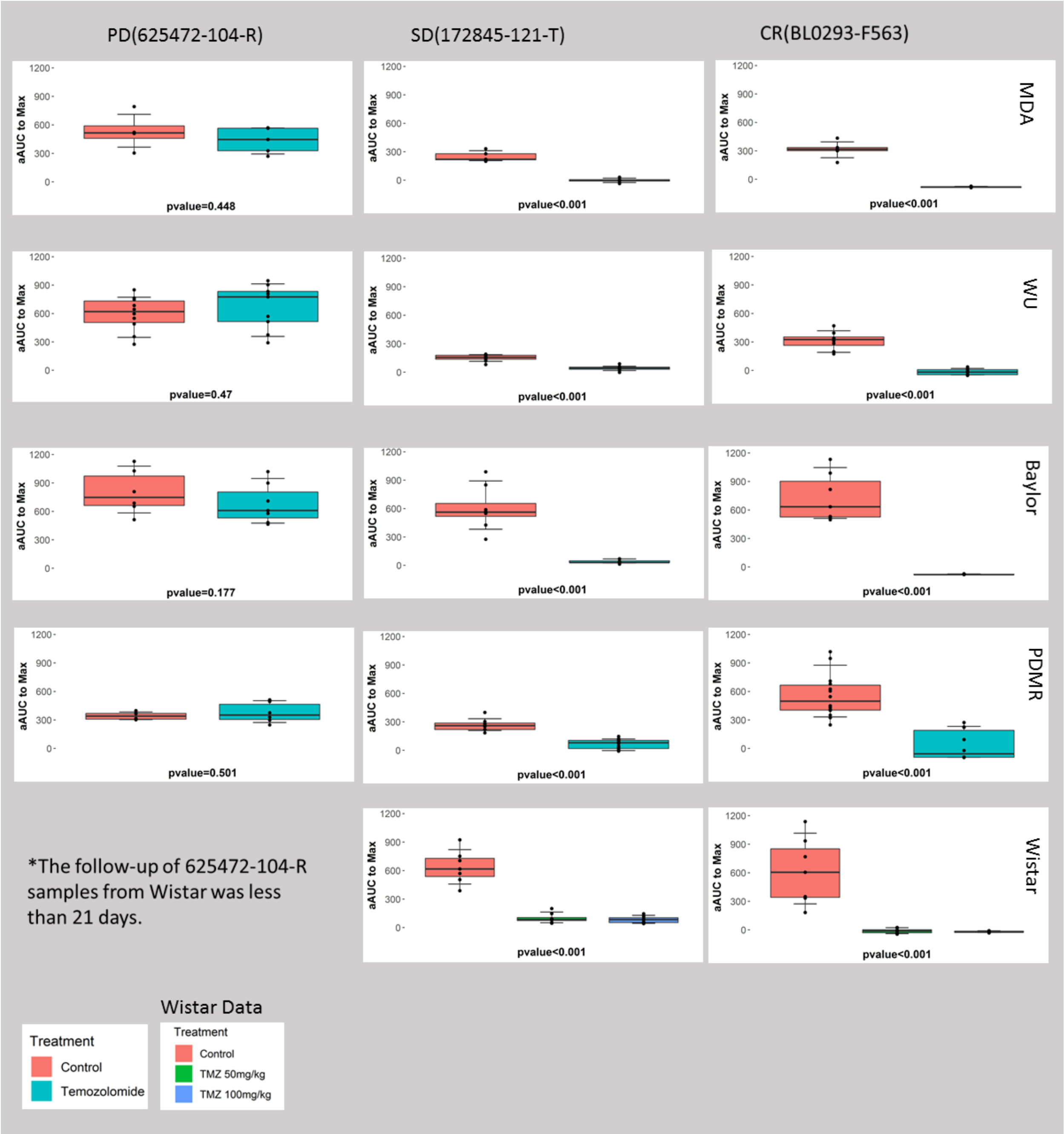

**Supplementary Figure 9.** Results, adjusted area under the curve to end date $\boldsymbol{aAUC_{max}}$. Analytical results of the adjusted area under the tumor growth curve from baseline to day of last measurement from various studies for progressive model (625472-104-R), stable disease model (172845-121-T) and complete response model (BL0293-F563) (columns 1-3, respectively).

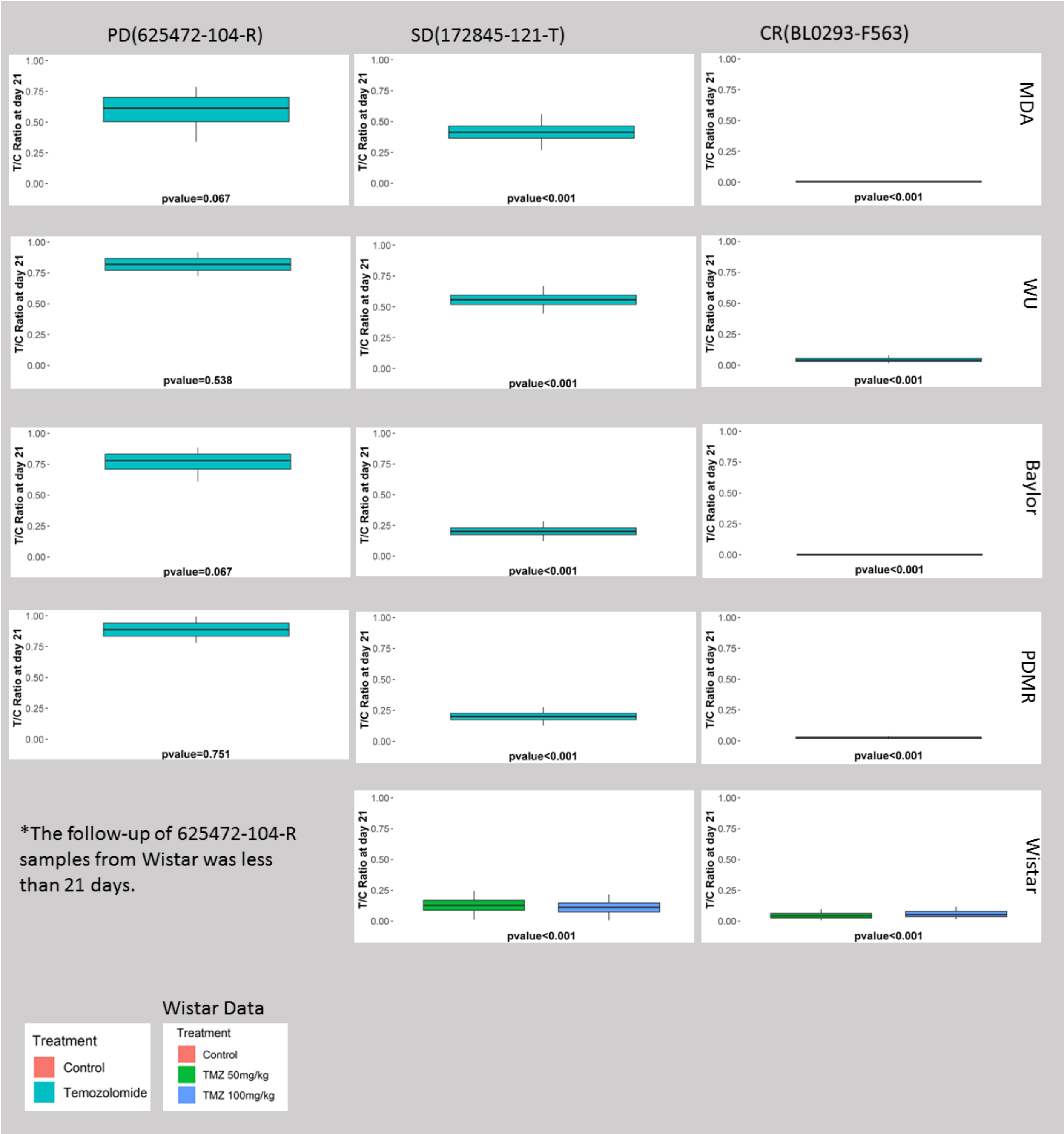

**Supplementary Figure 10.** Results, tumor growth inhibition to day 21 $TGI_{21}$. Analytical results of the tumor growth inhibition to day 21 measuring the mean ratio between treatment and baseline in terms of ratio of tumor size from baseline to day 21, from various studies for progressive model (625472-104-R), stable disease model (172845-121-T) and complete response model (BL0293-F563) (columns 1-3, respectively).

#### Supplementary Materials 3: Comparison of Tumor-normal whole exome workflow performance for each of the submitted workflows

The PDTCs submitted a total of five tumor-normal whole exome variant calling workflows. A summary of the tools used in each of the workflows is described in the Supplementary Table 2 below. Wiring schematics for each of the submitted workflows are shown in **Supplementary Figures 12 and 13.** Detailed command lines that were used to execute each workflow can be made available upon request. Detailed description about tools that each workflow comprises of can also be made available upon request via access to the workflow(s) on the CGC.

**Supplementary Table 2.** Main components of Tumor-Normal Whole Exome Workflows submitted by PDTCs

| **Workflows submitted by PDTCs** | **Main Components** |
|---|---|
| Workflow 1 | (BBSplit, BWA-MEM), **Manta**, **Strelka2**, USeq, Tabix |
| Workflow 2 | Trimmomatic, Xenome, BWA-alt-aware, **MuTect2**, snpEff, snpSift |
| Workflow 3 | combined human-mouse reference; BWA, samtools, **VarScan2** |
| Workflow 4 | BWA, custom mouse reads filtering; samtools, **MuTect2**, **VarScan2** |
| Workflow 5 | GATK, **VarScan2**, **Pindel**, **Strelka2**, samtools, VEP, snpSift, Picard |

**Supplementary Table 3.** Performance of the five PDTC workflows across SNPs at 10% and 50% mouse reads contamination.

| | Workflow 1<br>% | Workflow 2<br>% | Workflow 3<br>% | Workflow 4<br>% | Workflow 5<br>% |
|---|---|---|---|---|---|
| **SNP Precision** | | | | | |
| 10% Mouse | 96.4 | 94.0 | N/A | 2.2 | 56.5 |
| Contamination | 98.8 | 97.7 | 1.7 | 12.4 | 71.5 |
| | 99.0 | 98.5 | 39.1 | 13.3 | 75.5 |
| | 99.1 | 98.7 | 98.4 | 33.0 | 76.0 |
| | 99.1 | 98.7 | 99.1 | 45.0 | 76.0 |
| 50% Mouse | 96.3 | 94.0 | N/A | 2.2 | 56.3 |
| Contamination | 98.7 | 97.7 | 1.6 | 12.4 | 71.3 |
| | 99.0 | 98.5 | 38.3 | 13.3 | 75.3 |
| | 99.1 | 98.7 | 98.4 | 33.0 | 75.9 |
| | 99.1 | 98.7 | 99.1 | 45.0 | 75.8 |
| **SNP Recall** | | | | | |
| 10% Mouse | 23.9 | 20.7 | N/A | 2.6 | 31.7 |
| Contamination | 70.9 | 57.2 | 0.0 | 16.9 | 61.3 |
| | 91.3 | 87.4 | 0.5 | 18.4 | 75.2 |
| | 96.6 | 97.2 | 52.2 | 59.0 | 77.7 |
| | 97.3 | 98.6 | 97.2 | 97.9 | 78.0 |
| 50% Mouse | 23.9 | 20.7 | N/A | 2.6 | 31.7 |
| Contamination | 70.9 | 57.2 | 0.0 | 16.9 | 61.3 |
| | 91.3 | 87.4 | 0.5 | 18.4 | 75.2 |
| | 96.6 | 97.2 | 52.2 | 59.0 | 77.7 |
| | 97.3 | 98.6 | 97.2 | 97.9 | 78.0 |

a

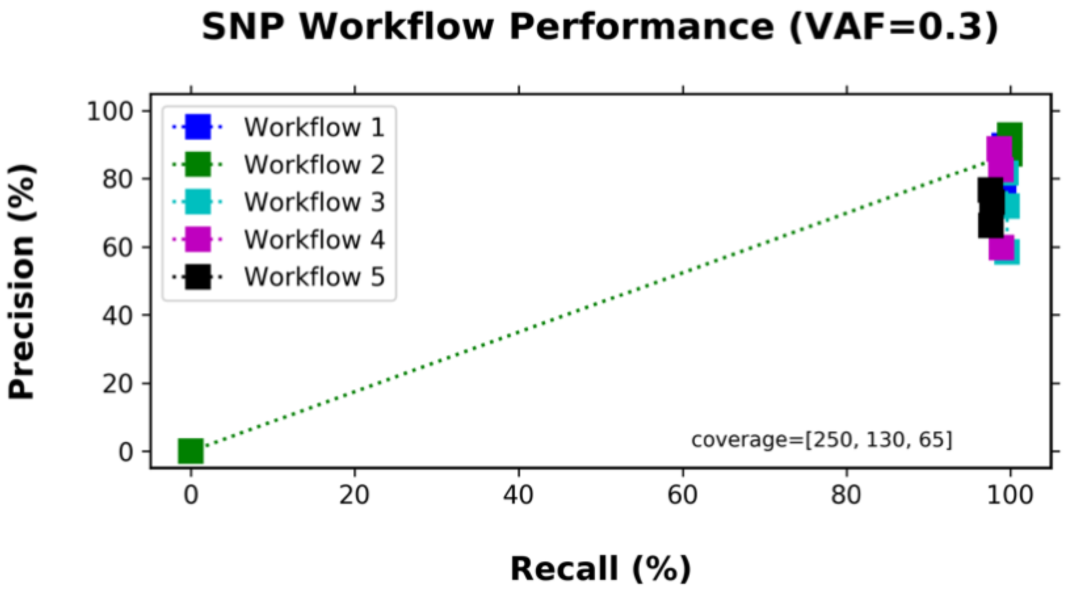

b

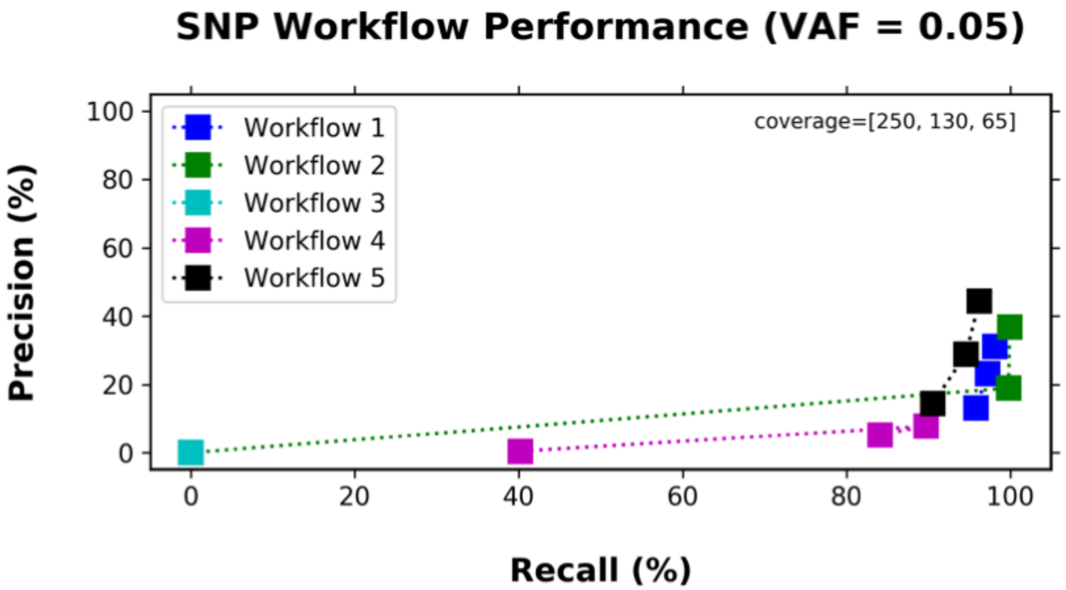

**Supplementary Figure 11:** Performance of the five PDTC workflows across SNPs at 250x, 130x and 65x coverage at 0.05 or 0.3 variant allele frequency (VAF). Precision and recall across SNPs of a simulated dataset (BS-DN) at 250, 130, and 65 coverage values and at two variant allele frequencies: **a)** 0.05 VAF and **b)** 0.3 VAF. As expected, workflow performance improved at higher VAFs and coverage. Workflow 2 that contains Mutect2 as the variant caller performs consistently the best across all samples in SNP calling at both 0.05 and 0.3 VAF.

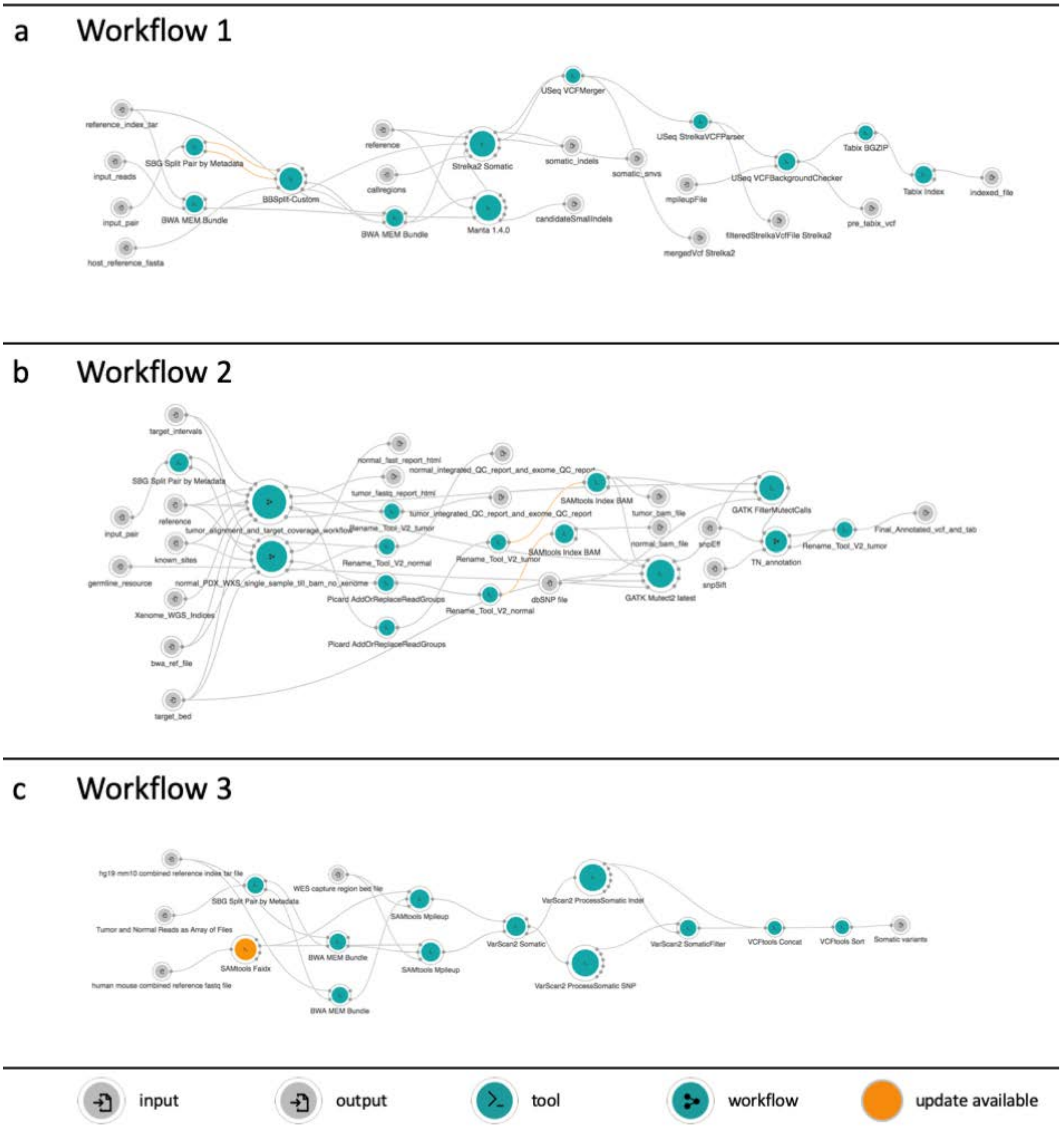

**Supplementary Figure 12:** Wiring diagrams for submitted whole exome workflows submitted by the PDX Development and Trials Centers. Wiring diagrams include nodes and connections. Nodes depict inputs - , outputs - , tools - , and workflows - . Connections between nodes depict that input to a node is from the output of another node. Orange nodes - identify a tool or a workflow with an available update. **a)** Workflow submitted by PDTC 1, **b)** Workflow submitted by PDTC 2, **c)** Workflow submitted by PDTC 3

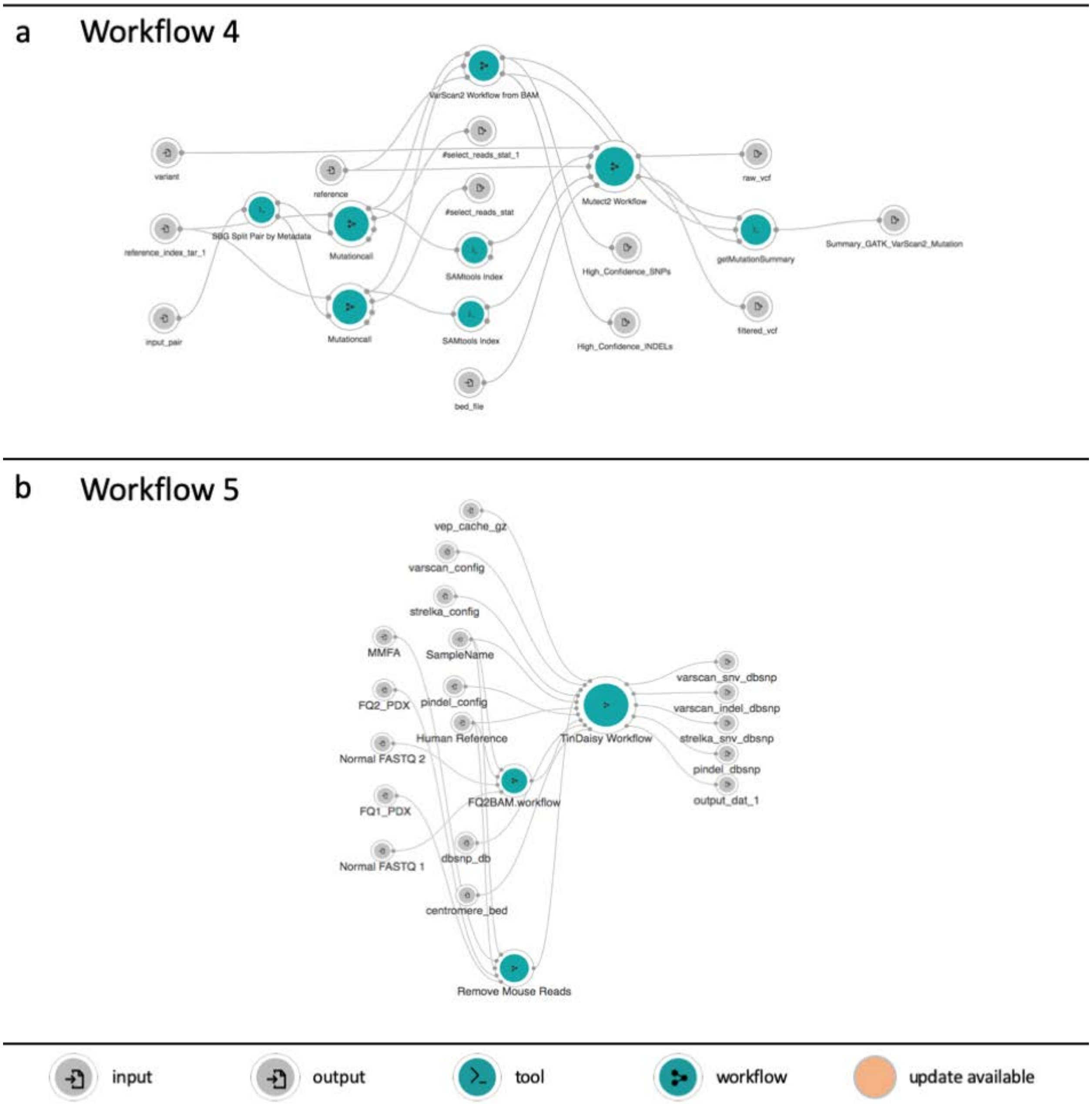

**Supplementary Figure 13:** Wiring diagrams for submitted whole exome workflows submitted by the PDX Development and Trials Centers. Wiring diagrams include nodes and connections. Nodes depict inputs - , outputs - , tools - , and workflows - . Connections between nodes depict that input to a node is from the output of another node. Orange nodes - identify a tool or a workflow with an available update. **a)** Workflow submitted by PDTC 4, **b)** Workflow submitted by PDTC 5

### Supplementary Materials 4: Workflows selected to process PDX whole exome and RNA-seq data

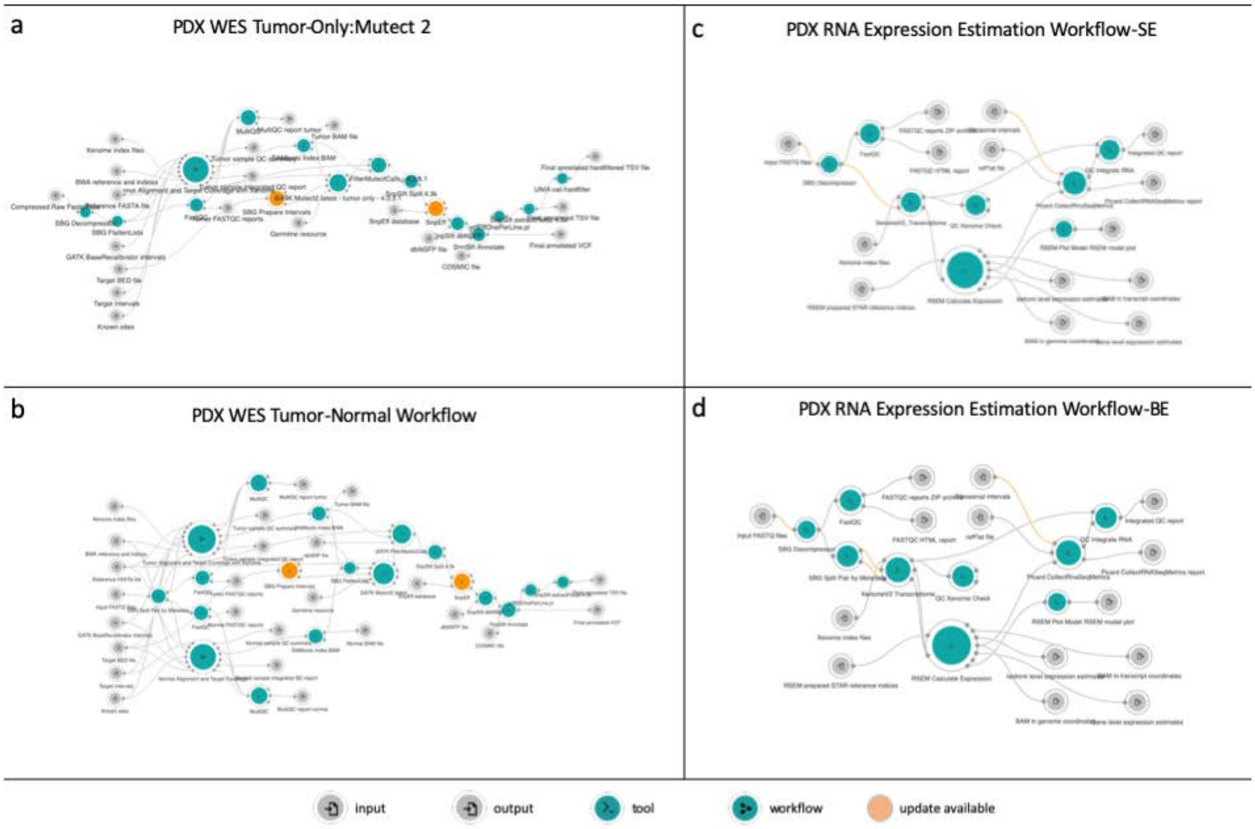

**Supplementary Figure 14: Workflow selected to process PDX Data.** The panel figure depicts whole exome and RNA-Seq workflows wiring diagram selected by the PDX Data Commons and Coordination Center (PDCCC) to process PDX data generated for this study. Wiring diagrams include nodes and connections. Nodes depict inputs - , outputs - , tools - , and workflows - . Connections between nodes depict that input to a node is from the output of another node. Orange nodes - identify a tool or a workflow with an available update. **a)** PDX whole exome tumor only workflow. The PDCCC selected the whole exome workflow after evaluating 5 workflows submitted by the PDX Development and Trial Center. **b)** PDX whole exome tumor-normal workflow. The PDCCC generated by the whole exome tumor-normal workflow by adapting the selected workflow described in a). **c)** PDX RNA Expression Estimation Workflow – Single Execution (SE). The PDCCC selected the RNA-seq workflow from established best practices. **d)** PDX RNA Expression Estimation Workflow – Batch Execution (BE). The PDCC adapted the PDX RNA Expression Estimation Workflow shown in c) to accept batch inputs.

### Supplementary Materials 5: Gene expression analysis using RNA-Seq workflow

#### Description

This RSEM workflow (RSEM 1.2.31) for quantifying gene expression uses the STAR aligner and is optimized to work with FASTQ input files.

To process multiple samples, please consider running batch tasks with this workflow and aggregating the results using Prepare Multisample Data workflow.

#### Essential Requirements

The following metadata fields are essential and should be assigned to input FASTQ files:

1. Sample ID: Any string. The identifier should be identical for both paired-end FASTQ files.
2. Paired-end: 1 or 2

This workflow will process both uncompressed and compressed FASTQ files (FASTQ.GZ, FASTQ.BZ2) and has been designed for paired-end data. By default, the workflow assumes unstranded data (Forward probability input parameter set to 0.5). Please adjust the value of this parameter (0.0 or 1.0) based on the library prep of your data.

The following output files will be generated:

- `Gene level expression estimates`
- `Isoform level expression estimates`
- `RSEM model plot`
- `BAM in transcript coordinates`
- `BAM in genome coordinates`
- `FASTQC reports ZIP archive`
- `FASTQC HTML report`
- `Integrated QC report`
- `Picard CollectRNASeqMetrics report`

#### Reference Files and Workflow Details

Required reference input files:

1. Xenome is used to classify reads as human or mouse. Xenome indices were built on hg38 and pseudoNOD transcriptome (based on SNP incorporation into mm10 genome from Sanger [ftp://ftp-mouse.sanger.ac.uk/REL-1505-SNPs_Indels/]). The default value of k=25 is used during the indices

preparation. Default file input: Xenome_transcriptome_indices_GRCh38_91_pseudoNod_mm10.tar.gz

2. STAR indices archive prepared by RSEM Prepare Reference (v.1.2.31). The default input file (GRCh38.91.chr_patch_hapl_scaf_rsem-1.2.31.star-index-archive.tar) was built using a GRCh38 FASTA file (primary assembly, EBV, alt contigs, decoys, and HLA contigs) and an annotation GRCh38 GTF file from Ensembl (release 91) (ftp://ftp.ensembl.org/pub/release-91/gtf/homo_sapiens/Homo_sapiens.GRCh38.91.chr_patch_hapl_scaff.gtf.gz)
3. refFlat file (hg38) used by Picard CollectRnaSeqMetrics tool. Downloaded from: http://hgdownload.cse.ucsc.edu/goldenPath/hg38/database/refFlat.txt.gz Default file input: refFlat.ucsc_hg38.txt
4. Ribosomal intervals (hg38) used by Picard CollectRnaSeqMetrics tool. Default file input: rRNA_hg38.interval

### Workflow Steps and Notable Parameters

#### Step 1: Optional input preprocessing

If FASTQ.BZ2 files are provided as inputs, the files will be decompressed before further analysis (as Xenome will only accept FASTQ and FASTQ.GZ files). FASTQ.GZ and uncompressed FASTQ input files will be passed on to other tools in the workflow.

#### Step 2: FASTQC analysis

Quality of the input FASTQ files is checked with FASTQC.

#### Step 3: Xenome classification of reads

FASTQ pairs are split (SBG Split Pair by Metadata) based on the appropriate paired_end metadata field values and classified by Xenome as mouse or human. QC Xenome Check tool checks that a sufficient number of reads have been classified as human. By default, minimum number of human reads required is set to 1000000, however this parameter is exposed (Minimum number of human-specific reads) and can be adjusted by the user. *Note*: If the Minimum number of human-specific reads cutoff is not met, the tasks will fail. If your expect <1000000 human reads in your input data, or are testing the workflow with subsetted files, please adjust this parameter accordingly.

#### Step 4: RSEM expression estimation

Expression is estimated using RSEM Calculate Expression tool (RSEM 1.2.31), with STAR as the aligner. Please ensure that the reference indices archive supplied to the tool has been prepared accordingly. RSEM Plot Model tool is used to generate RSEM plots.
Please note that by default, the workflow is setup to process unstranded data (Forward probability input parameter set to 0.5). Please make sure to adjust the value of this parameter (0.0, 0.5 or 1.0) based on the library-prep used.

#### Step 5: Additional QC

Additional QC reports are collected from Picard CollectRnaSeqMetrics tool and Xenome.

**Supplementary Table 4.** Total genomic region (in megabases [Mb]) covered by each capture

| Center | Capture Array Genome Coverage (Mb) |
|---|---|
| HCI-BCM | 115.23 |
| MD Anderson | 63.50 |
| Washington University | 39.11 |
| Wistar | 45.31 |

**Supplementary Table 5.** Mean number of calls after restricting to intersected array and applying allele frequency-based threshold

| Center | Average Number Variant Calls Center Specific Arrays | Average Number Variant Calls Intersected Arrays | Average Number Variant Calls AF Filtered Intersected Arrays |
|---|---|---|---|
| **HCI-BCM** | 77,885.33 | 5,333.00 | 3,286.00 |
| **MD Anderson** | 15,614.33 | 6,253.33 | 1,340.67 |
| **Washington University** | 14,569.67 | 8,103.67 | 3,274.00 |
| **Wistar Institute** | 11,618.00 | 5,406.00 | 3,114.67 |

**Supplementary Table 6.** RNA-Seq library-prep, sequencing type and depth for each sample

| Center | CaseID | PrepKit | Sequencing | total-reads |
|---|---|---|---|---|
| **HCI-BCM** | JAX-BL0293 | IlluminaTruSeqStandardRNAkitwithRiboZeroGold | Illumina HiSeq SE | 88,858,611 |
| | PDMR-172845 | IlluminaTruSeqStandardRNAkitwithRiboZeroGold | Illumina HiSeq SE | 83,090,055 |
| | PDMR-625472 | IlluminaTruSeqStandardRNAkitwithRiboZeroGold | Illumina HiSeq SE | 99,052,483 |
| **MDAnderson** | JAX-BL0293 | Kapa stranded mRNA-Seq Kit | Illumina HiSeq PE | 86,514,781 |
| | PDMR-172845 | Kapa stranded mRNA-Seq Kit | Illumina HiSeq PE | 71,158,865 |
| | PDMR-625472 | Kapa stranded mRNA-Seq Kit | Illumina HiSeq PE | 67,009,735 |
| **Washington University** | JAX-BL0293 | TruSeq Stranded Total RNA | NovaSeq S4 PE | 44,188,536 |
| | PDMR-172845 | TruSeq Stranded Total RNA | NovaSeq S4 PE | 56,610,898 |
| | PDMR-625472 | TruSeq Stranded Total RNA | NovaSeq S4 PE | 43,067,125 |
| **Wistar** | JAX-BL0293 | Lexogen mRNA | Illumina PE | 51,503,760 |
| | PDMR-172845 | Lexogen mRNA | Illumina PE | 53,648,665 |
| | PDMR-625472 | Lexogen mRNA | Illumina PE | 52,142,953 |

**Supplementary Table 7.** Sequencing depth and Mean Target coverage per sample across each center

| Center | CaseID | Total Reads | Mean Target Coverage |
|---|---|---|---|
| HIC-BCM | JAX-BL0293 | 84306497 | 182.489987 |
| | PDMR-172845 | 84500785 | 171.117572 |
| | PDMR-625472 | 69122847 | 144.39062 |
| MDAnderson | JAX-BL0293 | 24840423 | 34.378847 |
| | PDMR-172845 | 22722611 | 25.75321 |
| | PDMR-625472 | 23350341 | 28.91726 |
| Washington University | JAX-BL0293 | 63360889 | 225.221598 |
| | PDMR-172845 | 47171801 | 163.915966 |
| | PDMR-625472 | 36508439 | 131.624613 |
| Wistar | JAX-BL0293 | 249671426 | 192.639997 |
| | PDMR-172845 | 114733973 | 122.825594 |
| | PDMR-625472 | 129198866 | 133.174483 |

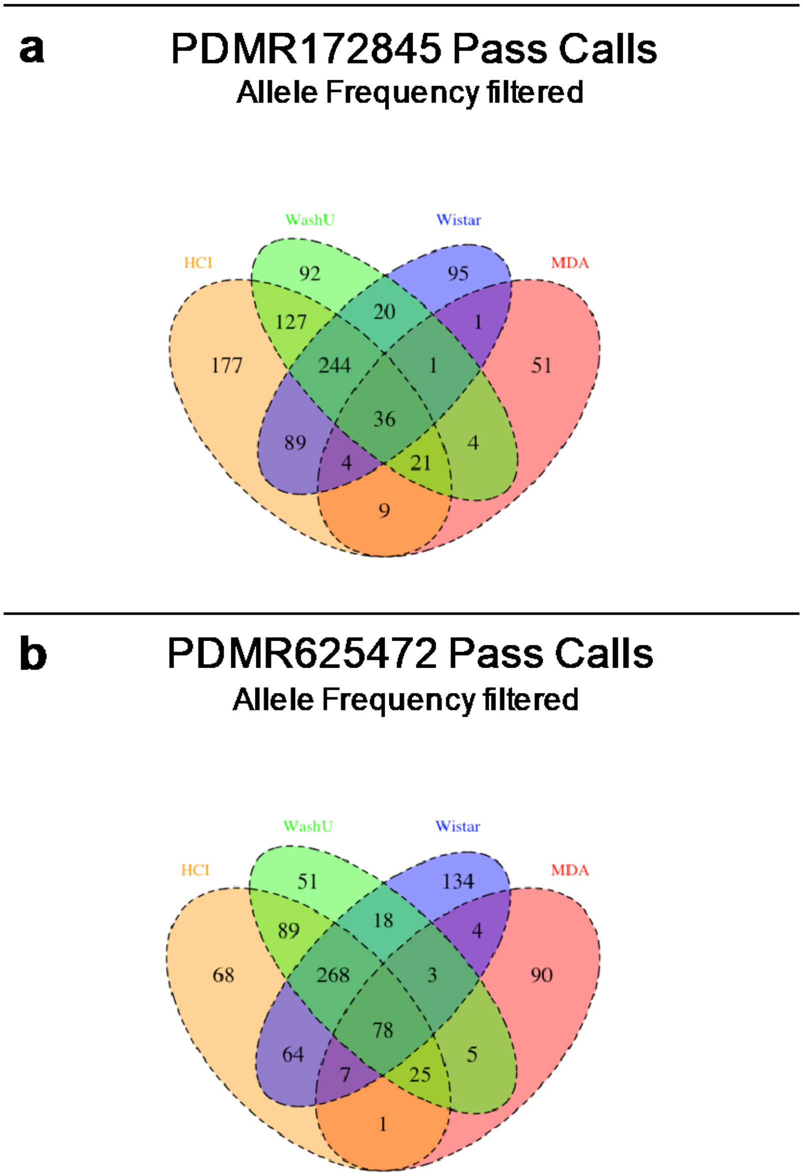

**Supplementary Figure 15.** A Venn diagram showing the overlap in high-quality variant calls among centers by model using intersected array and removing lower allele frequency (AF) calls. **a)** Pass variant calls for PDMR172845, **b)** Pass variant calls for PDMR625472.

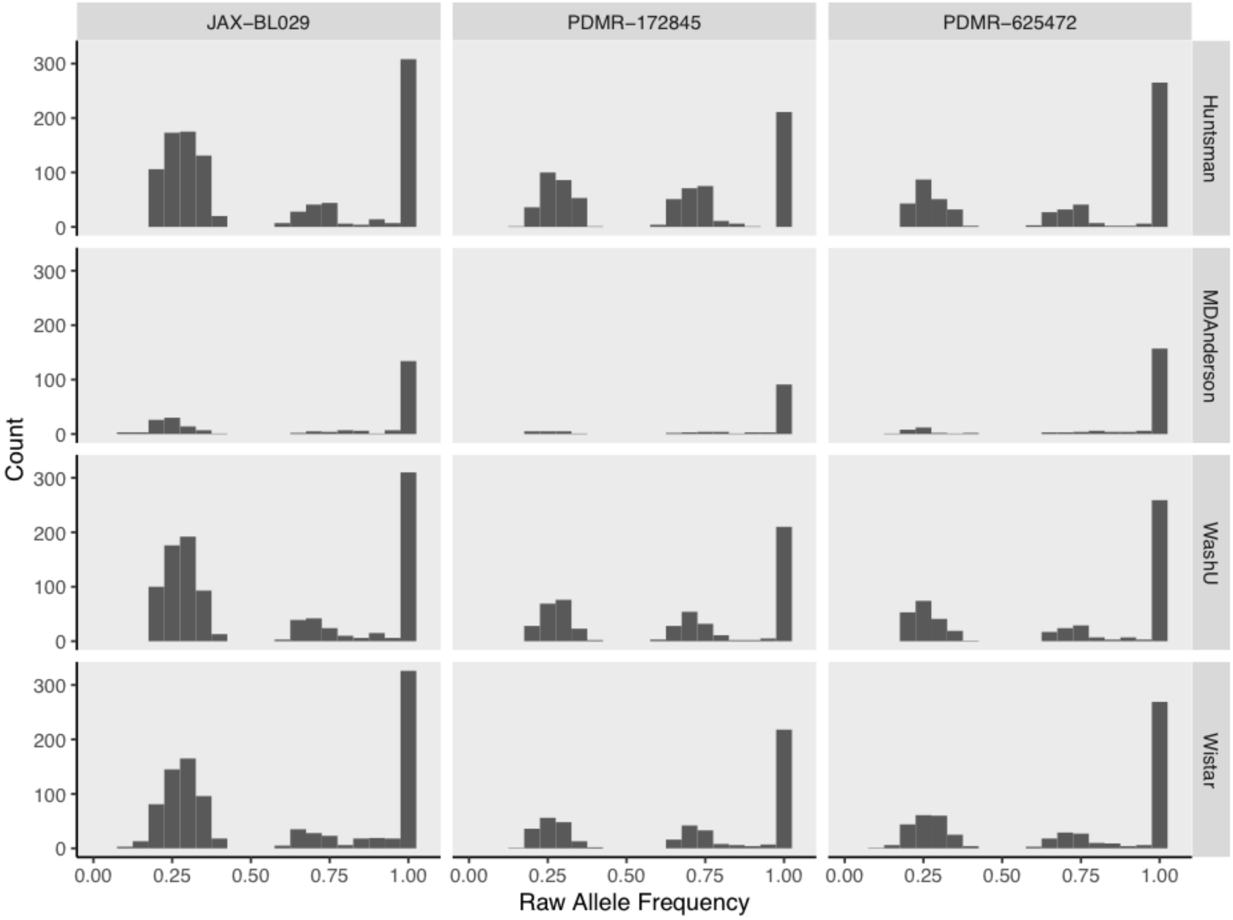

**Supplementary Figure 16.** Raw allele frequency for high-quality variants filtered to intersected array at allele frequency ≥ 0.2

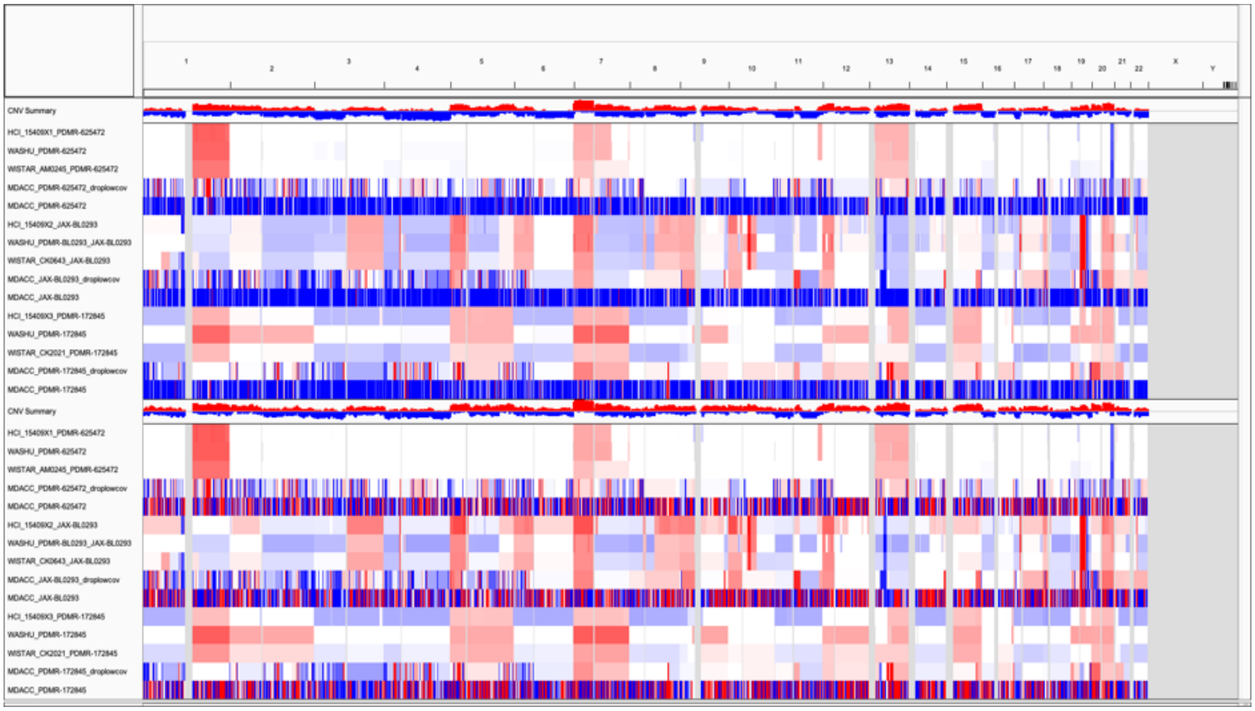

**Supplementary Figure 17.** An IGV snapshot of Median centered copy number alteration (CNA) segments

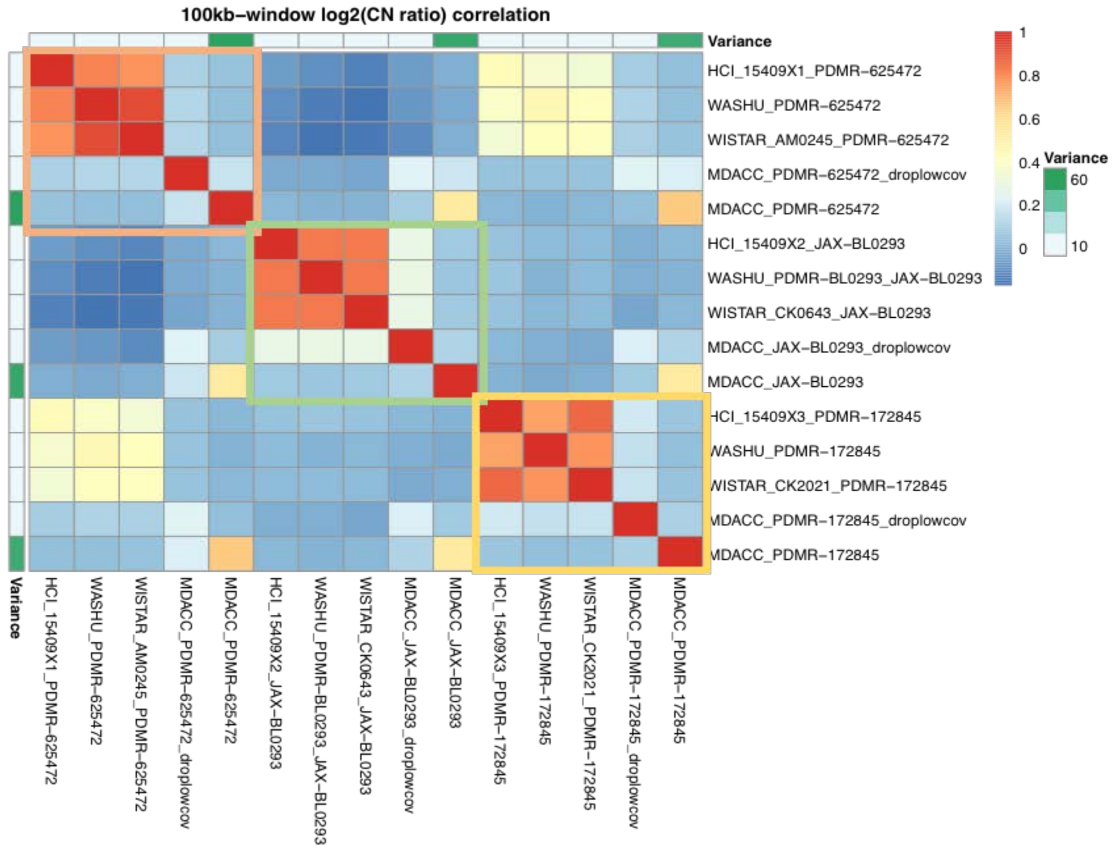

**Supplementary Figure 18.** A Pearson correlation coefficient based heatmap for the segmented log-2 copy number ratio across the 100kb-windows for each pair of samples

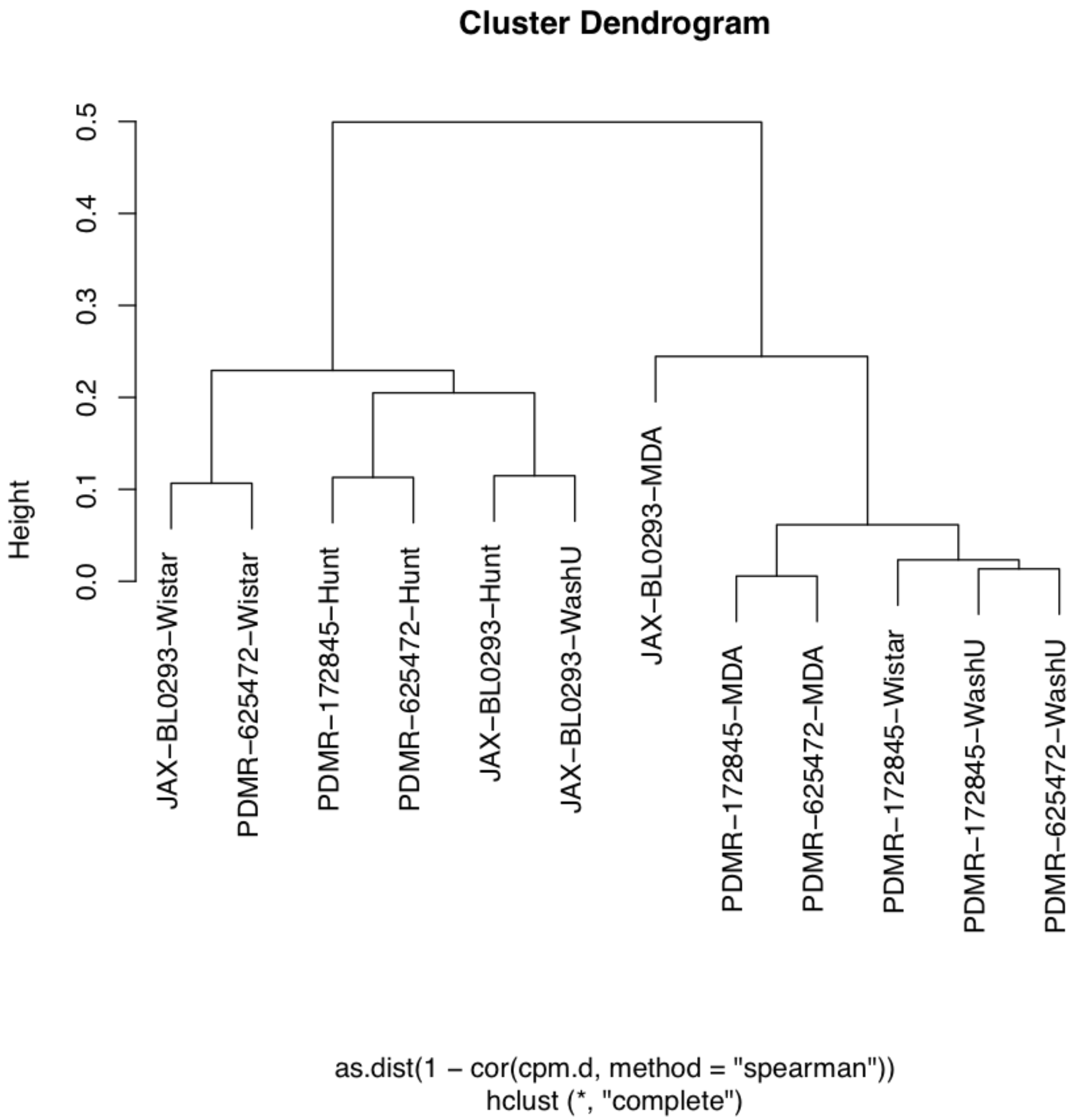

**Supplementary Figure 19.** Dendrogram of TMM normalized count per million values
